# Stress-dependent cell stiffening by tardigrade tolerance proteins through reversible formation of cytoskeleton-like filamentous network and gel-transition

**DOI:** 10.1101/2021.10.02.462891

**Authors:** Akihiro Tanaka, Tomomi Nakano, Kento Watanabe, Kazutoshi Masuda, Gen Honda, Shuichi Kamata, Reitaro Yasui, Hiroko Kozuka-Hata, Chiho Watanabe, Takumi Chinen, Daiju Kitagawa, Satoshi Sawai, Masaaki Oyama, Miho Yanagisawa, Takekazu Kunieda

## Abstract

Tardigrades are able to tolerate almost complete dehydration by entering a reversible ametabolic state called anhydrobiosis and resume their animation upon rehydration. Dehydrated tardigrades are exceptionally stable and withstand various physical extremes. Although trehalose and late embryogenesis abundant (LEA) proteins have been extensively studied as potent protectants against dehydration in other anhydrobiotic organisms, tardigrades produce high amounts of tardigrade-unique protective proteins. Cytoplasmic-abundant heat-soluble (CAHS) proteins are uniquely invented in the lineage of eutardigrades, a major class of the phylum Tardigrada and are essential for their anhydrobiotic survival. However, the precise mechanisms of their action in this protective role are not fully understood. In the present study, we first postulated the presence of tolerance proteins that form protective condensates via phase separation in a stress-dependent manner and searched for tardigrade proteins that reversibly form condensates upon dehydration-like stress. Through comprehensive analysis, we identified 336 such proteins, collectively dubbed “Desolvataion-induced ReversiblY condensing Proteins (DRYPs)”. Unexpectedly, we rediscovered CAHS proteins as highly enriched in DRYPs, 3 of which were major components of DRYPs. We revealed that these CAHS proteins reversibly polymerize into many cytoskeleton-like filaments depending on hyperosmotic stress in cultured cells and undergo reversible gel-transition *in vitro*. CAHS filamentation increases cell stiffness to resist deformation and improves resistance to dehydration-like stress. The conserved putative helical C-terminal region is necessary and sufficient for filament formation by CAHS proteins, and mutations disrupting the secondary structure of this region impaired both the filament formation and the gel transition. On the basis of these results, we propose that CAHS proteins are novel cytoskeletal proteins that form filamentous networks and undergo gel-transition in a stress-dependent manner to provide on-demand physical stabilization of cell integrity against deformative forces during dehydration and also contribute to the exceptional physical stability in a dehydrated state.

## Introduction

Water is an essential molecule for maintaining the metabolic activity and cellular integrity of living organisms. Some organisms, however, can tolerate almost complete dehydration by entering a reversible ametabolic state called anhydrobiosis [1]. Tardigrades, also known as water bears, are a prominent example of such desiccation-tolerant animals [2]. Under a drying environment, tardigrades gradually lose almost all body water and concurrently contract their bodies to a shrunken round form called a tun. Dehydrated tardigrades are exceptionally stable and can withstand various physically extreme environments including exposure to space [3,4]. Even after exposure to extreme stressors, tardigrades can reanimate within a few dozen minutes after rehydration.

Several tolerance molecules against dehydration stress have been identified in various organisms. One of the most analyzed molecules is the non-reducing disaccharide, trehalose. A significant amount of trehalose accumulates during desiccation in several anhydrobiotic animals, such as sleeping chironomids [5], brine shrimp [6], and some nematodes [7], some of which require trehalose synthesis for anhydrobiotic survival [8]. Trehalose is proposed to play its protective roles through 2 modes of action: water replacement, in which trehalose substitutes for water molecules; and vitrification, in which trehalose preserves cell components in an amorphous solid (glassy) state [9]. In tardigrades, however, no or only a little amount of trehalose accumulates, even in dehydrated states of the anhydrobiotic species [10], and a recent study suggested that trehalose synthesis genes in tardigrades are acquired in only limited lineages via horizontal transfer after the establishment of their anhydrobiotic ability [11], suggesting the presence of a trehalose-independent anhydrobiosis mechanism in tardigrades.

Late embryogenesis abundant (LEA) proteins are another example of tolerance molecules. LEA proteins are principally unstructured proteins originally identified in desiccating plant seeds and later found in several anhydrobiotic animals [12]. LEA proteins have many proposed roles, including stabilization of vitrified trehalose, molecular shielding of client biomolecules, and sequestration of ions [12]. LEA proteins can suppress dehydration-dependent denaturation of enzymes, and have strong synergistic protective effects with trehalose [13]. The LEA proteins of brine shrimp were recently reported to undergo phase separation to form droplet condensates upon dehydration and to increase the desiccation tolerance of insect cells [14].

Through a search for LEA-like heat-soluble proteins that remain soluble even after boiling in tardigrades, we previously identified cytoplasmic-abundant heat-soluble (CAHS) proteins from one of the toughest tardigrade species, *Ramazzottius varieornatus* [15]. CAHS proteins exhibited almost no similarity with non-tardigrade proteins, and later genome and transcriptome analyses suggested that CAHS proteins are present only in eutardigrades, one of the major classes of the phylum Tardigrada [11,16,17,18,19,20]. Despite the absence of sequence similarity between CAHS proteins and LEA proteins, they share similar biochemical properties, e.g., high-hydrophilicity supporting heat-solubility and structural transition from the disordered state in hydration to a helix under desolvating or dehydrated conditions [12,15]. Like LEA proteins, CAHS proteins can protect enzymes from dehydration stress [18] and *R. varieornatus* produces a remarkable amount of CAHS proteins rather than trehalose and LEA proteins. Knockdown of several CAHS genes that impaired the anhydrobiotic survival revealed that CAHS proteins are involved in the desiccation tolerance of eutardigrades [18]. Although CAHS proteins were proposed to act as a vitrifying agent based on a shift in differential scanning calorimetry, this hypothesis was recently counter-argued as such a shift could be explained by the evaporation of residual water [21], and the molecular mechanism remains to be elucidated.

Dehydration stress leads to the reduction of a cell volume and the destruction of cell structures, causing cells severe mechanical stress. To protect cells from these deformative forces, cytoskeletons like intermediate filaments (IFs) are generally principal players in counteracting mechanical stress in ordinary animal cells [22,23]. Interestingly, canonical cytoplasmic IFs were not found in Panarthropoda including tardigrades and arthropods. Tardigrades have a tardigrade-unique IF protein called cytotardin, which is not homologous to any cytoplasmic IFs in other animals and rather derives from the nuclear filament protein lamin [24]. Cytotardin does not localize to the nucleus because it lacks a nuclear localization signal, and instead forms belt-like filaments beneath the plasma membrane encircling epithelial cells, suggesting its contribution to the mechanical strengthening of epithelial tissues. In tardigrades, no IFs are known to form scaffold-like filamentous networks in the cytosol, which is thought to effectively counteract the deformative forces in many other animal cells [25,26].

In this study, we postulated the presence of tolerance proteins that form protective condensates in a stress-dependent manner, and comprehensively searched for such proteins in tardigrade lysate. Among more than 300 identified proteins that we collectively dubbed “desolvation-induced reversibly condensing proteins (DRYPs)”, we unexpectedly rediscovered CAHS proteins as highly-enriched and major components of DRYPs. Further analyses revealed that in response to stress, CAHS reversibly forms many cytoskeleton-like filaments in cultured cells and also exhibits reversible gelation *in vitro*. CAHS filamentation increases the mechanical strength of cultured cells and improves their resistance to dehydration-like stress. We also examined the structural basis required for filament formation by deletion and point mutation analyses. By studying the generated filament-defective mutants, we confirmed that the filament-forming ability is the basis for the gel transition of CAHS proteins. On the basis of these results, we propose a new tolerance model in which CAHS proteins act as a kind of cytoskeleton that reversibly forms intracellular filamentous networks in response to dehydration and induces gel transition that increases mechanical strength of cells and contributes to the desiccation tolerance of tardigrades.

## Results

### Desolvation-induced reversibly condensing proteins (DRYPs) are identified from *Ramazzottius varieornatus*

We designed the experimental scheme shown in Fig 1A to identify tardigrade proteins that form condensates in response to dehydration-like stress in a reversible manner. We began with the lysate of the desiccation-tolerant tardigrade species *R. varieornatus,* because this species constitutively expresses the tolerance proteins and its genome sequence is available [16]. First, we added trifluoroethanol (TFE) to a soluble fraction of *R. varieornatus* lysate to induce condensation in a dehydration-like state. TFE is a desolvating reagent that induces dehydration-like conformational changes in several desiccation-tolerance proteins, such as LEA and CAHS proteins [15,27,28]. The TFE-condensed proteins were collected as precipitates and resolubilized with TFE-free PBS to mimic rehydration. With 0% TFE treatment, no proteins were detected in the resolubilized fraction. In contrast, treatment with a higher concentration of TFE increased the number of proteins detected in the resolubilized fraction (Fig 1B and S1 Fig). As treatment with 20% and 30% TFE had similar effects, we considered 20% TFE to be an adequate stress condition for this screening (S1 Fig). When treated with TFE at 20% or higher, many proteins, especially those with a high molecular weight, were detected in the irreversibly precipitated fraction, indicating that only the selected proteins were retrieved in the resolubilized fraction.

**Fig 1.**
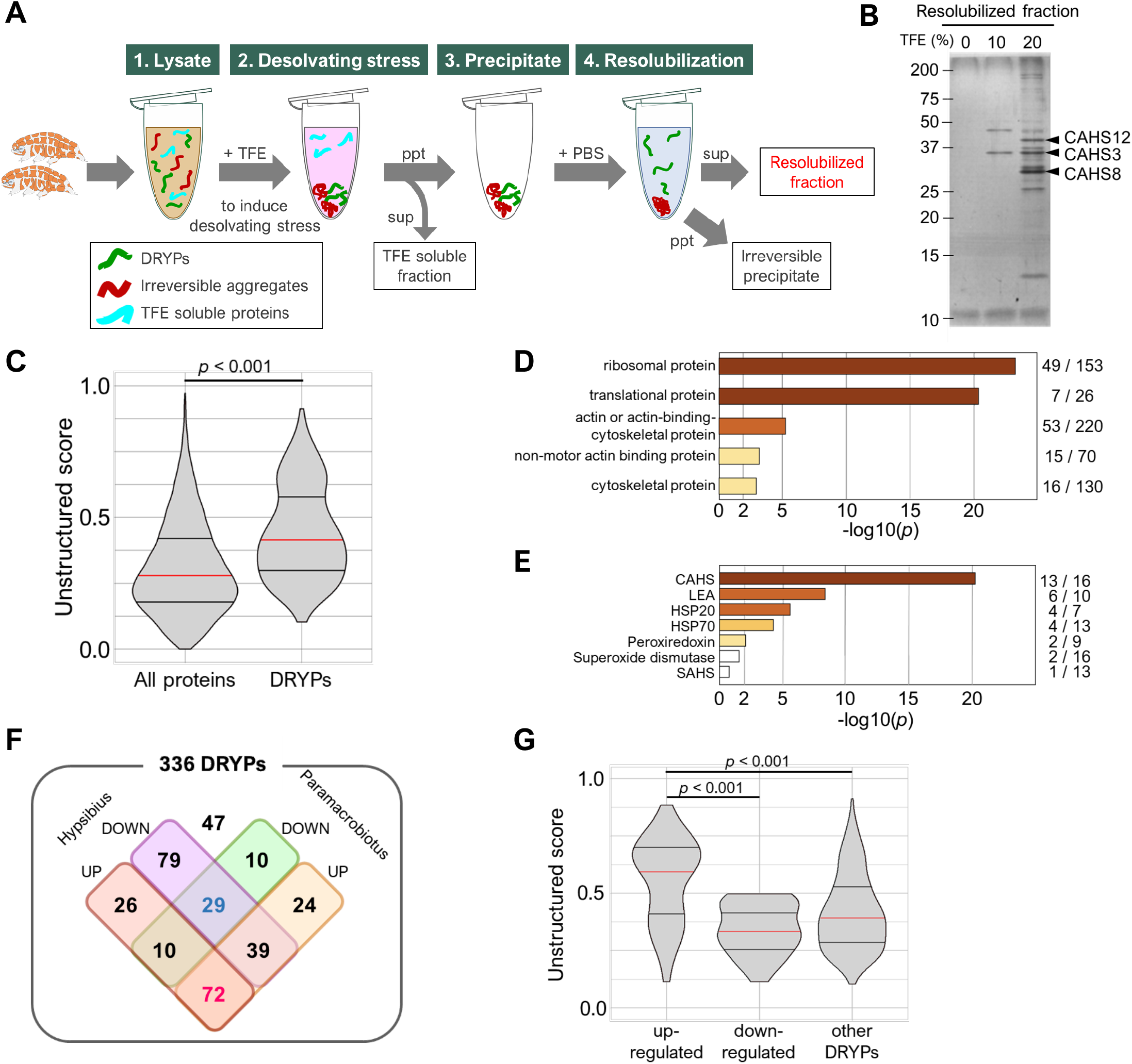
Isolation and characterization of desolvation-induced reversibly condensing proteins (DRYPs). (A) Experimental scheme of DRYP isolation from tardigrade lysate. (B) SDS-PAGE image of resolubilized fractions with 0%, 10%, or 20% TFE treatment. (C) Comparison of the unstructured-score distributions between all tardigrade proteins and DRYPs. (D) Enrichment analysis of the PANTHER protein class in DRYPs. Ribosomal proteins and cytoskeletal proteins were significantly enriched. The numbers of the corresponding proteins detected in DRYPs and all tardigrade proteomes are shown on the right, respectively. (E) Enrichment analysis of stress-related proteins in DRYPs. CAHS proteins were significantly enriched in DRYPs. (F) Venn diagram of DRYPs classified by up- or down-regulation upon desiccation in orthologs of 2 other tardigrade species. (G) Comparison of unstructured-score distributions among the differently regulated protein groups in DRYPs. “up-regulated” and “down-regulated” indicate up-regulated or down-regulated proteins in both species, respectively. Proteins up-regulated upon desiccation exhibited higher unstructured scores. Red and 2 black horizontal bars in violin plot indicate the 50th, 25th, and 75th percentiles, respectively. Statistical analyses were performed with the Wilcoxon rank sum test in (C) and the Steel-Dwass test in (G).

We identified 336 proteins in the resolubilized fraction (20% TFE) by liquid chromatography-tandem mass spectrometry (LC-MS/MS), and collectively termed these proteins “Desolvation-induced ReversiblY condensing Proteins (DRYPs)”. Reversible condensation is a characteristic property expected for unstructured proteins. To evaluate whether unstructured proteins are enriched in DRYPs, we calculated the unstructured score of each protein by IUPred2A and compared the score distribution between DRYPs and all tardigrade proteins. As expected, DRYPs contained a significantly higher proportion of unstructured proteins (p < 2.2e-16, Wilcoxon rank sum test; Fig 1C). We assigned *Drosophila melanogaster* orthologs for tardigrade proteins and performed enrichment analysis of PANTHER Protein class or Gene Ontology term in DRYPs. The results revealed that ribosomal proteins and actin-related cytoskeletal proteins were well concentrated in DRYPs (Fig 1D and S2 Fig). Among DRYPs, however, 105 (31%) proteins had no apparent fly orthologs and DRYPs contain many tardigrade-unique proteins (21%) including known tolerance proteins like CAHS proteins. Therefore, we expanded the enrichment analyses to the previously annotated tardigrade tolerance protein families that contain more than 5 members [16], and revealed the significant enrichment of CAHS, LEA, HSP20, HSP70 and peroxiredoxin families in DRYPs (p < 0.01, chi-square test; Fig 1E), suggesting that our new screening scheme concentrates desiccation-tolerance related proteins to the resolubilized fraction. To evaluate this possibility further, we classified DRYPs into 3 groups: stress-upregulated groups, stress-downregulated groups, and the others. *R. varieornatus* is one of the toughest tardigrade species that constitutively expresses stress-related genes [16]. Thus, we utilized gene expression data of 2 closely related tardigrades, *Hypsibius exemplaris* and *Paramacrobiotus* sp. TYO, both of which exhibit strong up-regulation of tolerance gene expression upon desiccation [11,17]. Of 336 DRYPs, 315 proteins had orthologs in both species and 72 genes were upregulated during dehydration (Fig 1F). Statistical analysis indicated that the up-regulated proteins were significantly enriched in DRYPs compared with the tardigrade proteome (p = 9.53e-29, chi-square test). In addition, the up-regulated proteins also exhibited a much higher unstructured score (Fig 1G), suggesting that tolerance-related unstructured proteins were well concentrated in the resolubilized fraction in our scheme. Because CAHS proteins were highly enriched in the DRYPs (Fig 1E), and also 3 major bands in the resolubilized fraction were separately identified as CAHS12, CAHS3, and CAHS8 (Fig 1B and S3 Fig), we focused on these 3 CAHS proteins for further analyses.

### CAHS3, CAHS8, and CAHS12 reversibly assemble into filaments or granules in animal cells depending on hyperosmotic stress

To visualize the stress-dependent condensation, 3 CAHS proteins, such as CAHS3, CAHS8 and CAHS12 proteins were separately expressed as a GFP-fused protein in human cultured HEp-2 cells and the distribution changes of these fusion proteins were examined upon exposure to a hyperosmotic stress, which induces water efflux like dehydration stress [29]. In an unstressed condition, CAHS3-GFP broadly distributed in the cytosol, whereas CAHS8-GFP and CAHS12-GFP distributed broadly in both the cytosol and the nucleus with CAHS12-GFP showing a slight preference for the nucleus (Fig 2A). When exposed to hyperosmotic medium supplemented with 0.4 M trehalose, CAHS3-GFP condensed and formed a filamentous network in the cytosol (Fig 2A and 2B). Similar filament formation was observed when CAHS3 alone was expressed without GFP (S4 Fig), suggesting that filament formation is an intrinsic feature of CAHS3 protein rather than artifact of fusion with GFP. CAHS12-GFP also formed filaments in the cytosol and more prominently in the nucleus in a majority of cells, though granule-like condensates were also observed in the nucleus of approximately 34% of the cells (Fig 2B and S5 Fig). CAHS8-GFP predominantly formed granule-like condensates especially in the nucleus, but filaments were also observed in the cytosol in a small population (~ 3%) of the cells. Similar distribution changes were observed even when GFP was fused to the opposite site in CAHS proteins (S6 Fig), while GFP alone did not exhibit such drastic changes. When hyperosmotic stress was removed by replacing with isosmotic medium, all CAHS condensates, both filaments and granules, rapidly dispersed (Fig 2A and 2B). Hyperosmotic stress by other supplemented osmolytes, such as 0.2 M NaCl or 0.4 M sorbitol, which have an equivalent osmolarity to 0.4 M trehalose, induces similar filament or granule formation, suggesting that the condensations of CAHS proteins are induced by hyperosmotic stress itself rather than specific effects of each osmolyte (S7 Fig). Similar reversible condensations of CAHS proteins were also observed when expressed in *Drosophila* cultured S2 cells (S8 Fig and S1 Movie), indicating that the stress-dependent filament/granule condensations are intrinsic features of CAHS proteins commonly observed in animal cells of taxonomically distant species.

**Fig 2.**
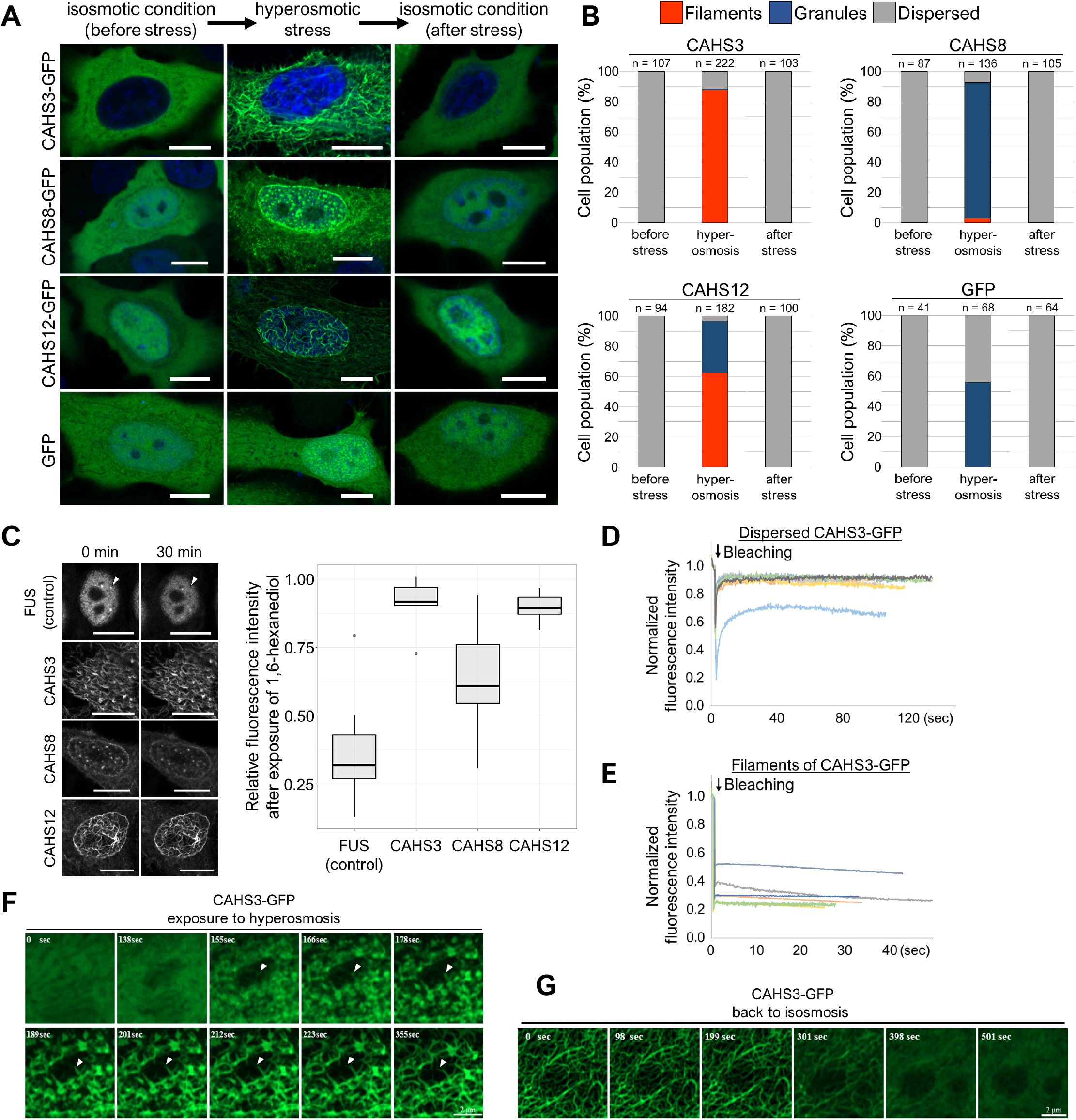
Reversible formation of filaments or granules by CAHS3, CAHS8, and CAHS12 proteins in response to a hyperosmotic stress. (A) Distribution changes in AcGFP1-tagged CAHS3, CAHS8, or CAHS12 proteins in HEp-2 cells during the transient hyperosmotic treatment in which the cells were exposed to HBSS containing 0.4 M trehalose. Blue indicates Hoechst33342 staining of nuclei. Scale bar, 10 μm. (B) The proportion of distribution patterns (filaments, granules, or dispersed) of each CAHS protein in human cells. (C) Effects of the liquid droplet disruptor, 1,6-hexanediol on condensates of FUS (n = 15), CAHS3 (n = 7), CAHS8 (n = 24), and CAHS12 (n = 7). FUS is a control protein sensitive to 1,6-hexanediol. Box plots show the distributions of the fluorescence intensity at 30 min relative to that at 0 min. Center bar and edges indicate 50th, 25th, and 75th percentiles, respectively and whiskers correspond to the 1.5 interquartile range. Scale bar, 10 μm. (D and E) Fluorescence recovery after photobleaching (FRAP) analyses of CAHS3-GFP in human cells in dispersed state under an isosmotic condition (D, n = 7), and in a filament-formed state under a hyperosmotic condition (E, n = 6). (F and G) Time-lapse images of filament formation or deformation of CAHS3-GFP in human cells (see also S2 and S3 Movies). (F) CAHS3-GFP first condensed into granules (155 s) and then elongated into filaments (355 s) as indicated by white arrows. (G) CAHS3-GFP filaments simultaneously collapsed and dispersed (398 s). Time since the medium exchange to hyperosmotic (F) or isosmotic (G) solution is shown in each image. Scale bar, 2 μm.

Granule-like condensates of CAHS8 resemble droplet structures formed by intrinsically disordered proteins via liquid-liquid phase separation. To test this possibility, we examined the effect of 1,6-hexanediol, a disruption reagent of liquid-like condensates. After treatment with 5% 1,6-hexanediol for 30 min, the well-known droplet-forming protein FUS effectively dispersed, while several CAHS8 granules in the nucleus also dispersed but much less effectively than FUS protein granules (Fig 2C), suggesting that CAHS8 granules were partly liquid-like, i.e., between liquid and solid states. In contrast, the filament structures of CAHS3 or CAHS12 were not affected by the hexanediol treatment, suggesting that CAHS3 and CAHS12 filaments were in a static solid-like state.

To further assess the staticity of CAHS filaments, we performed fluorescence recovery after photobleaching (FRAP) analysis on CAHS3-GFP both before and after exposure to hyperosmotic stress. In unstressed cultured cells, CAHS3-GFP was broadly distributed in the cytosol and the bleached fluorescence was rapidly recovered (Fig 2D), indicating their high mobility nature. In contrast, under hyperosmotic stress, CAHS3-GFP filaments exhibited almost no fluorescence recovery after bleaching (Fig 2E). These results demonstrated that CAHS3 molecules freely disperse in an unstressed condition, but upon the exposure to hyperosmotic stress, CAHS3 molecules are firmly integrated into a filament and lose their mobility.

To elucidate the process of filament formation and deformation in more detail, we captured time-lapse images of cells expressing CAHS3-GFP while changing the stress conditions by high-speed super-resolution microscopy. Approximately 2.5 min after the medium was changed to a hyperosmotic condition by a perfusion device, CAHS3-GFP began to condense simultaneously at many sites in the cells and rapidly formed fibril structures. The fibrils then further extended in a few dozen seconds (Fig 2F and S2 Movie). When the hyperosmotic stress was removed by changing to an isosmotic medium, CAHS3 filaments simultaneously began to loosen and gradually dispersed in approximately 6 min (Fig 2G and S3 Movie). The initial condensation of CAHS3 and the granule formation of CAHS8 likely occurred via phase-separation, which frequently leads to co-condensation of multiple proteins, especially those containing similar motifs [30]. CAHS proteins share several conserved motifs and could thus cooperatively form the same condensates. To examine this, we co-expressed pairs of the 3 CAHS proteins labeled with different fluorescent proteins in human cells. Under hyperosmotic stress, CAHS3 filaments did not co-localize with CAHS8 granules or CAHS12 filaments (Fig 3), suggesting no interaction between them. In contrast, CAHS8 largely co-localized with CAHS12 filaments throughout the cell, suggesting that the granule-forming CAHS8 cooperatively forms the filament structure with other CAHS proteins such as CAHS12.

**Fig 3.**
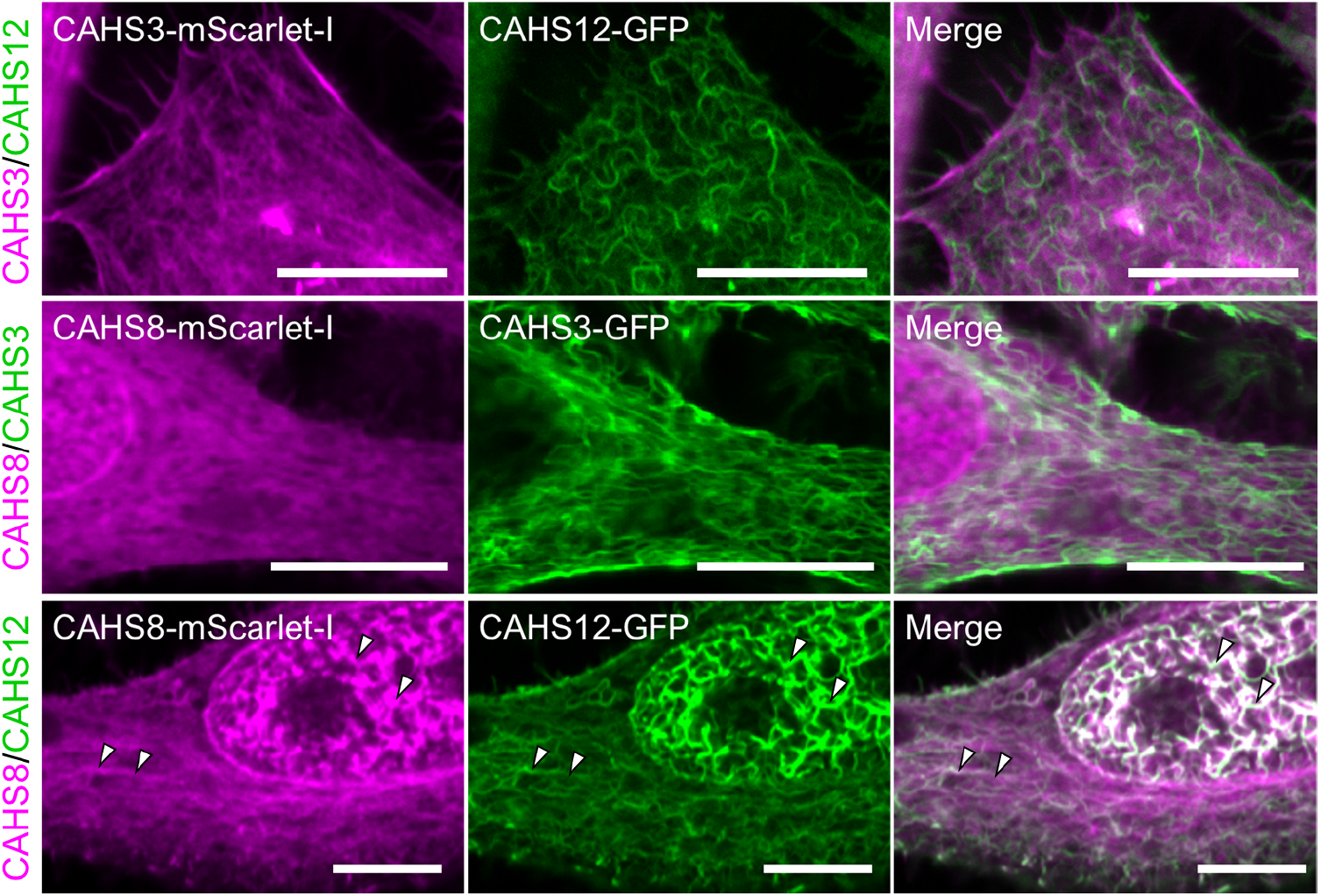
Cooperative filament formation of CAHS8 with CAHS12. Fluorescent images of HEp-2 cells co-expressing pairs of CAHS3, CAHS8, and CAHS12 proteins with a different fluorescent-tag under a hyperosmotic condition. CAHS3 co-localized with neither CAHS8 nor CAHS12. In contrast, CAHS8 well co-localized with CAHS12 filaments. White arrowheads indicate representative co-localization. Scale bar, 10 μm.

### Filament formation of CAHS3 or CAHS12 is independent of other cytoskeletons

Filamentous networks formed by CAHS3 or CAHS12 proteins resembled cytoskeletal structures. To examine whether the CAHS proteins formed filaments exclusively or cooperatively with other cytoskeletal structures or organelles, we co-visualized major cytoskeletons or organelles by expressing cytoskeleton/organelle marker proteins tagged with fluorescent proteins and then compared those with the distribution of CAHS-GFP filaments in human cells. As shown in Fig 4A–4C and S9A–S9D Fig, the CAHS3-GFP filaments and CAHS12-GFP filaments did not overlap with any examined cytoskeleton and organelles, such as microtubules, various intermediate filaments, mitochondria, and endoplasmic reticulum, except for a slight co-localization with actin filaments. Because GFP alone also exhibited partial co-localization with actin filaments under a hyperosmotic condition (Fig 4D), we assumed that the GFP-moiety is responsible for this slight co-localization between CAHS-GFP and actin filaments. To clarify the independence of CAHS filament formation from actin filaments, we treated the cells with an actin polymerization inhibitor, cytochalasin B. As a result, actin-filaments were significantly disrupted, but CAHS filaments were not affected (Fig 4E), suggesting that filament formation of both CAHS3 and CAHS12 is independent from actin filaments. CAHS8-GFP unexpectedly co-localized with an intermediate filament, vimentin (S9B Fig), in addition to actin filaments. CAHS8 could interact with vimentin filaments under hyperosmotic stress in human cells, though no vimentin genes were found in the tardigrade genome [24].

**Fig 4.**
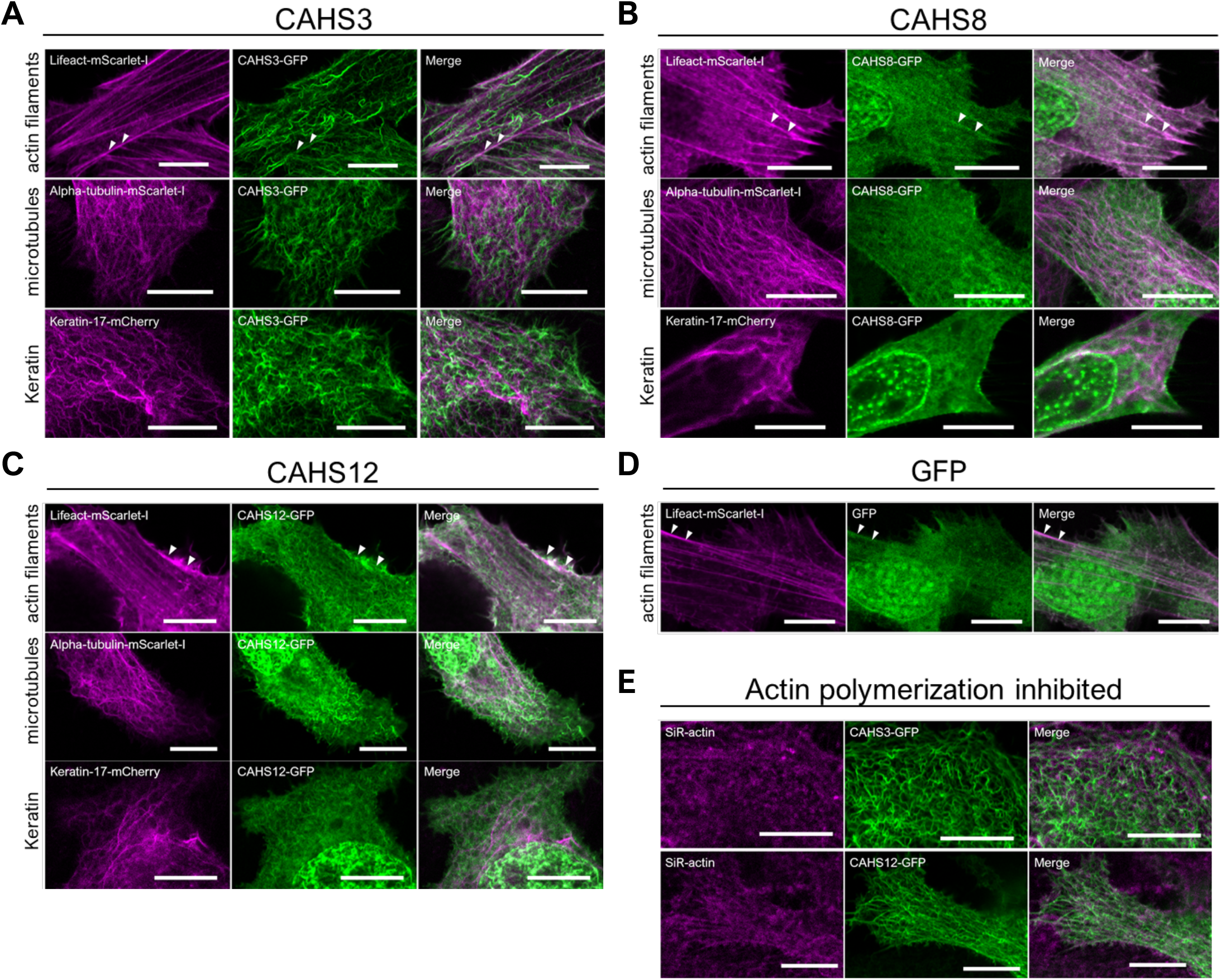
CAHS filaments are independent structures of other cytoskeletons. (A–C) Confocal images of AcGFP1-tagged CAHS proteins and fluorescently labeled cytoskeletal proteins in HEp-2 cells under a hyperosmotic condition. White arrows indicate slight co-localization of CAHS proteins and actin filaments. (D) Co-localization analysis of GFP alone and actin filaments. GFP alone partly co-localized with actin filament under a hyperosmotic condition. (E) Effects of the actin polymerization inhibitor cytochalasin B on CAHS filaments. Depolymerization of actin filaments had no effect on the formation of CAHS filaments. Scale bar, 10 μm.

### C-terminal regions are necessary and sufficient for filament-formation by both CAHS3 and CAHS12

To reveal the structural basis of CAHS filament formation, we first searched and found 10 conserved motifs by comparing 40 CAHS proteins of 3 tardigrade species, *R. varieornatus, H. exemplaris,* and *Paramacrobiotus* sp. TYO (S10 and S11 Figs). In particular, we found that 2 C-terminal motifs (CR1 and CR2) are highly conserved in all CAHS family members except 1 CAHS protein of *H. exemplaris* (Fig 5A and S10 Fig). To determine the region responsible for filament formation, we generated a series of truncated mutant proteins of CAHS3 or CAHS12 either N-terminally or C-terminally, and examined their filament formation in human cultured cells under a hyperosmotic stress (Fig 5B and 5C). In CAHS3, N-terminal deletion to motif 3 or C-terminal deletion to CR2 drastically impaired filament formation and instead granule formation was frequently observed in the cytosol (Fig 5B and S12 Fig). Accordingly, we designed a truncated mutant consisting of the minimum required region from motif 3 to CR2 (motif 3-motif H1-CR1-CR2), and revealed that this region is sufficient for the filament formation by CAHS3 protein (Fig 5B). Similarly, in CAHS12 protein, the region consisting of CR1, CR2, and the 2 preceding motifs (motif H2-motif H3-CR1-CR2) was shown to be necessary and sufficient for the filament formation (Fig 5C). These results indicated that 2 highly conserved motifs (CR1 and CR2) and 2 preceding motifs (65~85 residues) play an essential role in the filament-formation of both CAHS3 and CAHS12 proteins.

**Fig 5.**
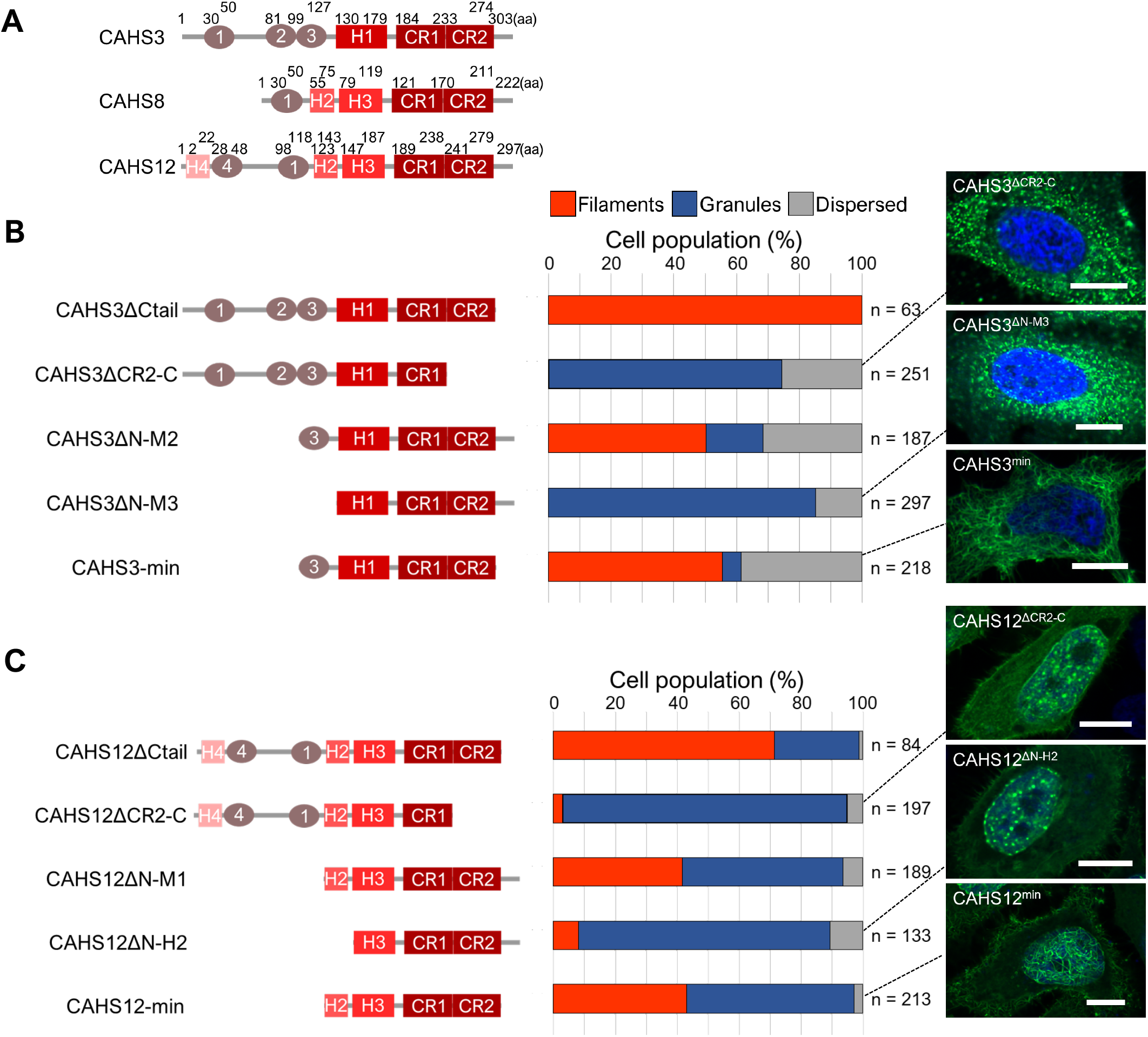
Conserved C-terminal regions are necessary and sufficient for the filament formation of CAHS3 and CAHS12. (A) Schematic diagrams of CAHS3, CAHS8 and CAHS12 proteins. “CR1” and “CR2” indicate putative helical motifs highly conserved among almost all CAHS family members. “H1”, “H2”, “H3”, and “H4” indicate putative helical conserved motifs. “1”, “2”, “3”, and “4” indicate other conserved motifs. (B and C) Schematic diagrams and the corresponding distribution patterns of the truncated mutants of CAHS3 (B) or CAHS12 (C). Quantified cell proportions of the distribution patterns under a hyperosmotic condition are shown as a stacked bar graph. Confocal images are shown for the representative distribution pattern of the corresponding CAHS mutants. Blue indicates Hoechst33342 staining of nuclei. Scale bar, 10 μm.

### Helix-disrupting mutations in CR impair filament formation of CAHS3 and CAHS12

In the regions responsible for the filament formation of both CAHS3 and CAHS12 proteins, extensive helix and putative coiled-coil structures were predicted by the secondary structure prediction tool, JPred4 and COILS (Fig 6 and S13 Fig). The coiled-coil structure is the key structural basis for the polymerization of intermediate filaments [31]. To test whether these predicted secondary structures are important for filament formation, we generated 2 mutants for each CAHS3 and CAHS12 by substituting leucine with proline, which are predicted to disrupt the helical and coiled-coil structures of CR1 or CR2, respectively (Fig 6) [32]. As expected, all coiled-coil disruption mutants failed to form filaments and instead formed granules (Fig 6 and, S14 and S15 Figs). The double mutation (CAHS3-L207P-L236P) further suppressed filaments formation and even reduced granule formation (Fig 6A and S16 Fig). These results suggested that the secondary structures of both CR1 and CR2 are an important basis for the filament formation by CAHS3 and CAHS12 proteins.

**Fig 6.**
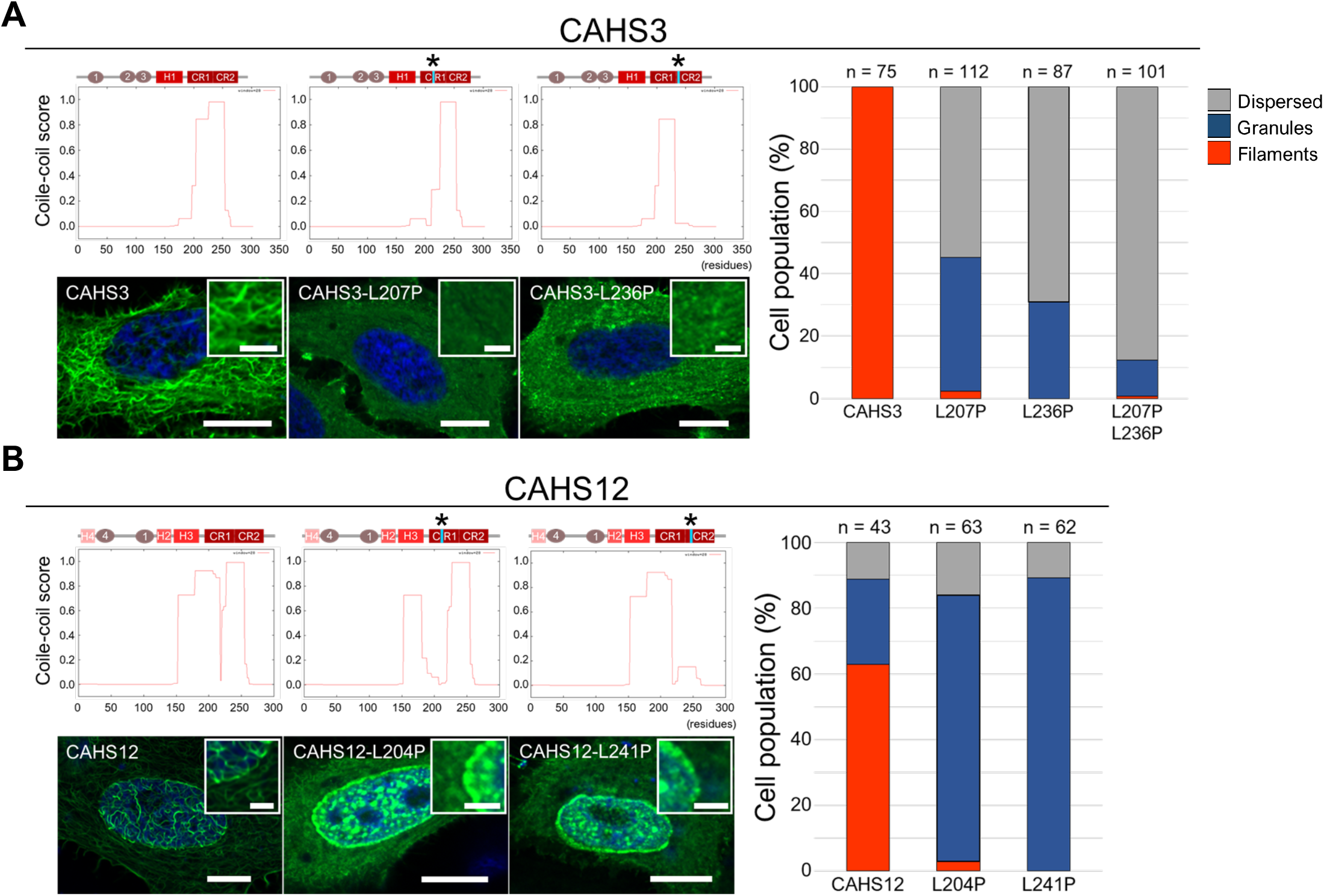
Suppression of filament-formation by mutations disrupting the coiled-coil structure in the conserved region of CAHS3 and CAHS12. (A and B) Effects of a helix-disrupting mutation by substituting leucine with proline on filament formation are shown for CAHS3 (A) and CAHS12 (B). Schematic structure and the coiled-coil score predicted by COILS are shown for both wild-type and proline substitution mutants. Asterisks indicate the sites of proline substitutions. Substitution with proline substantially decreased the coiled-coil score in the corresponding region. Confocal images show representative distribution patterns of the corresponding CAHS proteins (Scale bar, 10 μm). Enlarged image is shown as superimposition in each panel (Scale bar, 2.5 μm). Blue indicates Hoechst33342 staining of nuclei. Quantified cell proportions of each distribution pattern are shown as stacked bar plots on the right.

### *In vitro* reversible gel transition of CAHS proteins depending on desolvating reagent and salt

To examine whether CAHS proteins alone are sufficient to form filaments, we performed *in vitro* experiments using purified CAHS3-GFP proteins. Under an unstressed condition, the uniform distribution of CAHS3-GFP proteins was observed under a confocal microscope (Fig 7A). When the desolvating reagent TFE was added to induce a dehydration-like conformational change as in our initial screening, CAHS3-GFP immediately condensed and formed mesh-like fibril networks after 1 min. This result indicated that CAHS3 proteins alone can sense the changes in the condition and form filaments without the assistance of other proteins.

**Fig 7.**
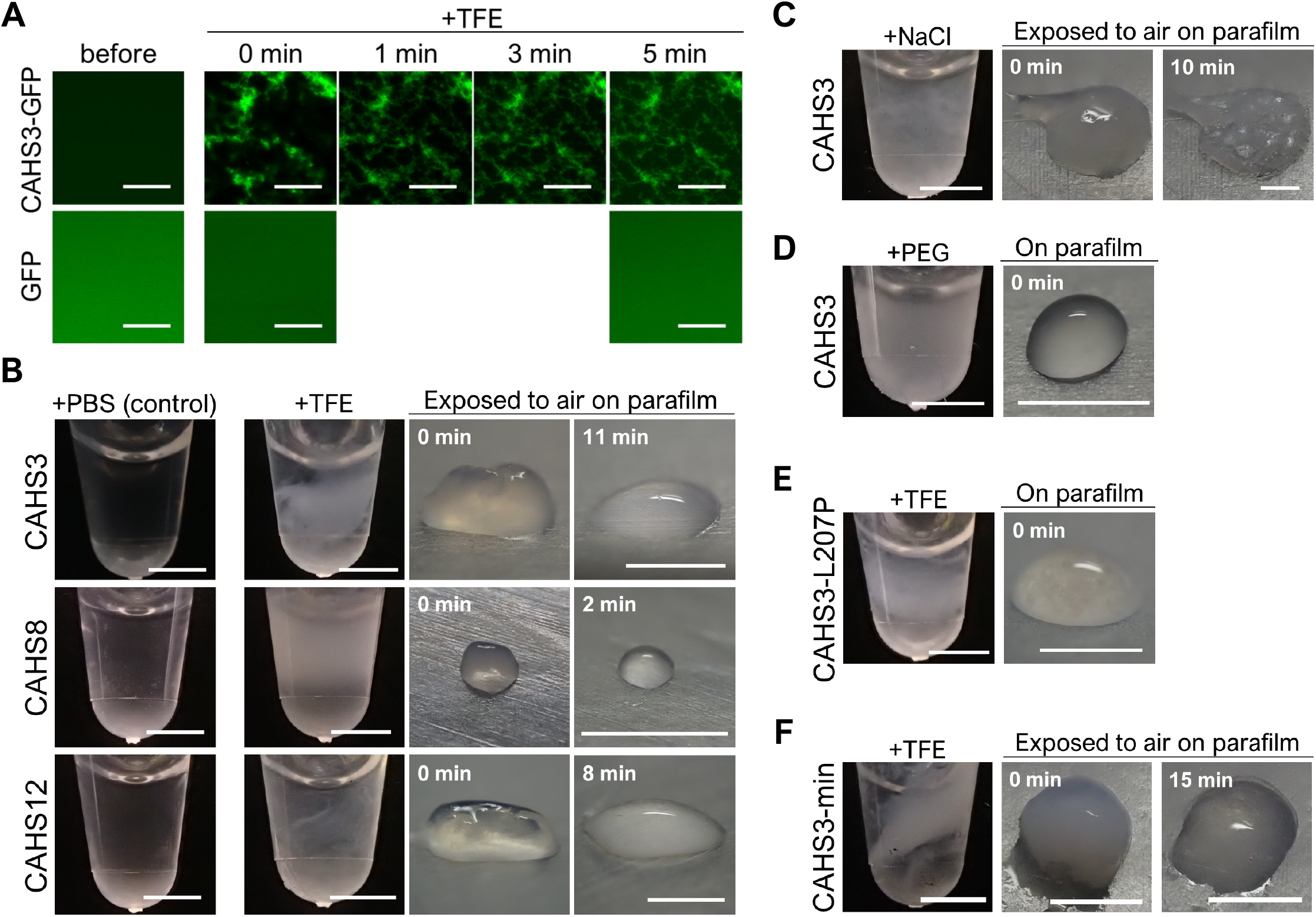
Gel transition of CAHS proteins upon desolvating or salt stress *in vitro*. (A) *In vitro* timelapse confocal images of fibril formation of CAHS3-GFP proteins (1.24 mg/mL) after adding TFE (final 20%). GFP is a non-filament forming control. (B) TFE-dependent reversible gel-formation of CAHS proteins. By adding TFE (final 20%), CAHS3, CAHS8, and CAHS12 protein solutions (4.0 mg/mL) became turbid and transited into a gel-like state. The gels spontaneously liquefied within several minutes (shown in white letters) after exposure to air. (C) Persisting gelation of CAHS3 induced by the addition of NaCl (2 M). (D) Addition of the molecular crowding agent, polyethylene glycol (PEG, final 20%) induced turbidity, but no gelation. (E) Filament-defective CAHS3-L207P mutant protein solutions failed to transit into a gel-like state under 20% TFE. (F) Minimum filament-forming CAHS3 truncated protein (CAHS3-min) solution reversibly solidified under 20% TFE like full-length CAHS3 protein. Scale bar, 20 μm in (A), 2 mm in (B–F)

When TFE was added to the solution containing a higher concentration of purified CAHS3 protein (final 4 mg/mL; S17 Fig), the protein solution immediately became turbid, and the solution was solidified into a gel-like state (Fig 7B, upper panels). When the CAHS3 gel in the tube was spread onto parafilm, the CAHS3 gel spontaneously liquefied within approximately 10 min. We speculated that volatilization of TFE relieved the desolvating stress, thereby making the CAHS3 gel resoluble. Consistently, washing with TFE-free PBS also redissolved the gelated CAHS3 (S18 Fig). While the control protein BSA was not solidified in the same condition (S19 Fig), CAHS8 and CAHS12 exhibited a similar TFE-dependent reversible gel-transition like CAHS3, but the gel of CAHS8 was much smaller than those of other CAHS proteins (Fig 7B, middle and lower panels), suggesting differences in the propensity for gelation among CAHS proteins. We also examined whether other stressors that could emerge during dehydration induce CAHS gelation and revealed that an increased concentration of salt (2 M NaCl) also induced the gel-transition of CAHS3 proteins (Fig 7C), while a molecular crowding agent (20% polyethylene glycol) caused turbidity, but no gelation (Fig 7D). The salt-induced gel persisted even after exposure to air on parafilm, possibly because salt cannot evaporate (Fig 7C). The granule-forming CAHS8 only formed a very small gel *in vitro,* implying a possible relationship between the filament-forming ability in cells and the gel-forming ability *in vitro*. This notion was supported by the fact that the filament-deficient CAHS3-L207P mutant protein failed to form the gel *in vitro* (Fig 7E). In contrast, minimum CAHS3 protein possessing the filament-forming ability (CAHS3-min) successfully formed the gel *in vitro* upon the addition of TFE and this transition was reversible as in full-length CAHS3 (Fig 7F), suggesting that filament-forming ability underlies the gel transition of CAHS proteins *in vitro*.

### Gelation of CAHS3 improves mechanical strength of a cell-like microdroplet

To reveal what the gelation of CAHS proteins provides, we evaluated the effects of CAHS gelation on the mechanical properties of cells using cell-like microdroplets covered with a lipid layer. The elasticity of the microdroplets was examined by measuring the elongation length in a micropipette while aspirating with a certain pressure. Microdroplets containing uniformly distributed CAHS3-GFP exhibited continuous elongation exceeding 50 μm under very small pressure (<< 0.5 kPa), indicating that they were not elastic and in a liquid phase (Fig 8). On the other hand, the addition of salt induced the filament formation by CAHS3-GFP and the corresponding microdroplets exhibited significant elasticity (Young’s modulus ~2.0 kPa in average), indicating that the CAHS3-GFP droplets gelated and then physically hardened. Microdroplets containing GFP alone were not elastic regardless of the addition of salt (Fig 8B and 8C).

**Fig 8.**
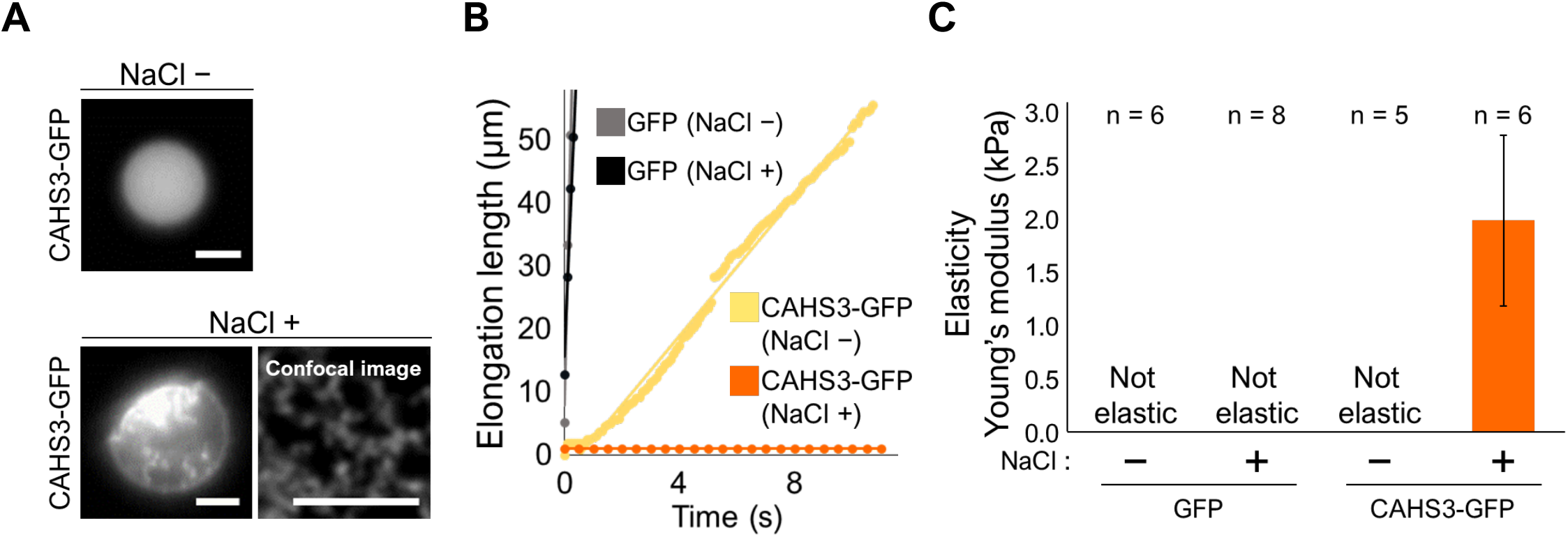
CAHS gelation increases the mechanical strength of cell-like microdroplets. (A) Representative fluorescent images of a microdroplet containing CAHS3-GFP in the absence or presence of additional NaCl. Scale bar, 5 μm. (B) Representative response curves of the elongation length of microdroplets containing CAHS3-GFP or GFP alone under a very small pressure (<< 0.5 kPa). Continuous elongation exceeding 50 μm indicates not elastic and in a liquid phase. (C) Comparison of the elasticity (Young’s modulus) among droplets containing CAHS3-GFP or GFP with or without NaCl addition. Data are presented as average ± SE.

### CAHS3 confers mechanical resistance against deformation by hyperosmotic stress on insect cells

To further determine the effects of CAHS filament formation on animal cells, we established a *Drosophila* S2 cell line stably expressing CAHS3. S2 cells lack canonical cytoplasmic intermediate filaments as tardigrade cells do [24] and thus it would be suitable to measure the effect of CAHS filamentation. As CAHS3 filamentation increased the mechanical strength of microdroplets *in vitro*, we also examined the effect of CAHS3 filamentation on the elasticity of the S2 cells by measuring the cortical cell stiffness using an atomic force microscope (AFM). Under an unstressed condition, the CAHS3-expressing cells exhibited no significant difference in the elasticity with that of the control cells transfected with empty vector. Under a hyperosmotic condition, control cells exhibited higher elasticity than that in an unstressed condition, but the CAHS3-expressing cells exhibited significantly higher elasticity than that of the control cells under the same condition (p < 0.05; Fig 9A), which is consistent with the results using microdroplets. Hyperosmotic stress reduces the cell volume through osmotic pressures [29]. As CAHS3-expressing cells exhibited higher elasticity under a hyperosmotic condition, they somewhat counteract the pressure through forming cytoskeletal-filamentous network and might resist the cell shrinkage. To examine this possibility, we measured the cell volume changes after exposure to hyperosmotic stress. As shown in Fig 9B, CAHS3-expressing cells retained the cell volume significantly better than the control cells (p < 0.001; Fig 9B). These results suggest that CAHS filament formation stiffen cells and protect them from deformation stress caused by dehydration-like stress. Finally, we examined the effect of CAHS3 on cell viability after exposure to hyperosmotic stress for 48 h. Cell viability was evaluated by the exclusion of propidium iodide (PI), which is an indicator of cell integrity. Under an unstressed condition, CAHS expression did not affect the cell viability, but after 48 h treatment with hyperosmotic stress, CAHS3-expressing cells exhibited the increased cell viability (Fig 9C and D). The stabilization of cell structure by CAHS proteins may contribute to the survival of tardigrade cells during the dehydration process.

**Fig 9.**
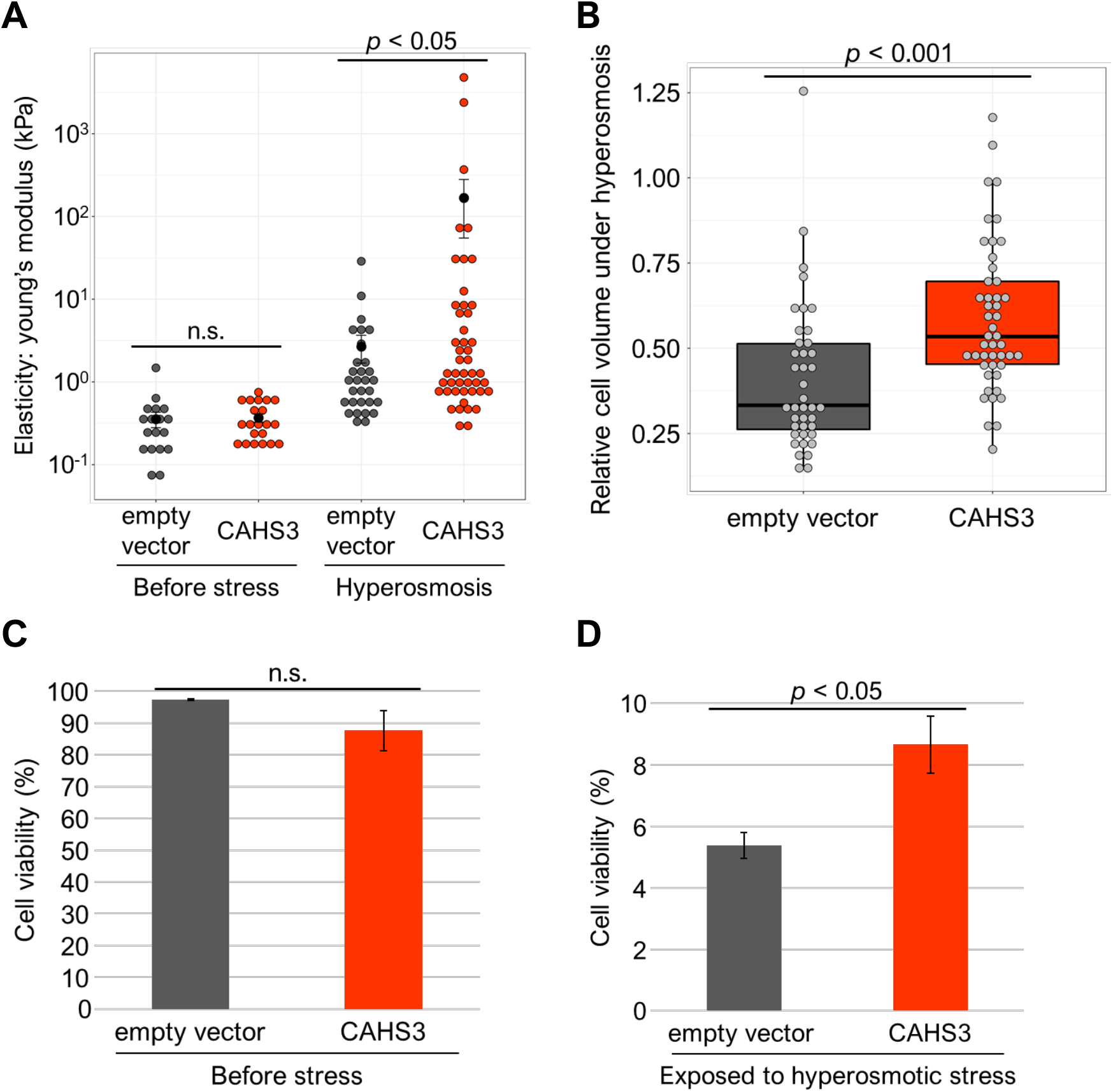
CAHS-expressing cells exhibit higher resistance to deformation under hyperosmotic stress. (A) The effects of CAHS3-filamentation on the cortical elasticity of *Drosophila* S2 cells. CAHS3-stably expressing cells exhibited higher elasticity compared to the control cells transfected with empty vector under a hyperosmotic condition supplemented with 0.4 M trehalose for 3 h. Gray and red dots indicate the values of each measurement. Black dots and bars indicate averages and standard errors, respectively. (B) Comparison of cell volume changes by hyperosmotic stress between CAHS3-expressing cells and control cells. The relative cell volume was calculated by dividing the volume under hyperosmotic stress by the averaged cell volume under isosmotic conditions. Center bar and edges indicate 50th, 25th, and 75th percentiles, respectively and whiskers correspond to the 1.5 interquartile range. (C and D) Comparison of cell viability between CAHS3-expressing cells and control cells under an isosmotic condition (C) and a hyperosmotic condition for 48 h (D). Propidium iodide was used to determine dead cells. Survival rates were examined in 6 wells for each condition by counting > 500 cells/well. Statistical analyses were performed with the Wilcoxon rank sum test in (A) and (B), and Student’s t-test in (C) and (D). n.s. means not significant in the statistical tests.

## Discussion

Our study provides evidence that CAHS proteins reversibly condense in a stress-dependent manner and form a cytoskeleton-like filamentous network in animal cells or undergo gel-transition *in vitro* (Figs 2A and 7B), and we further demonstrated that the CAHS filamentation increase the mechanical strength of cell-like microdroplets and improve the resistance to deformation stress of insect cells (Figs 8 and 9). In the previous study, CAHS proteins were suggested to act as a vitrifying agent like trehalose during dehydration based on the shift in differential scanning calorimetry (DSC) [18], but this hypothesis was recently counter-argued with data demonstrating that the shift in DSC can be explained by water retention of CAHS proteins [21]. Because hydrogel generally has high water retention properties, our observation of gel-transition by CAHS proteins supports the water retention in the counterargument rather than vitrification. *In vitro* gel transition was observed when using a relatively high concentration (~ 4 mg/mL) of CAHS protein solution (Fig 7B), and the filament-defective CAHS mutants failed in transition to gel (Fig 7E), suggesting that a dense filament formation is the structural basis for the gel transition of CAHS proteins. To confirm the protein concentration used in gel-transition *in vitro* is physiologically relevant, we estimated the amount of endogenous CAHS3 proteins in *R. varieornatus* by immunoblotting analysis, indicating that the amount of CAHS3 proteins is about 3.8 ng per individual (S20 Fig; see Materials and Methods). The wet weight of a single individual of *R. varieornatus* was reported to be 1.84 μg [33] which roughly corresponds 1.84 nL, and thus our rough estimate of the concentration of endogenous CAHS3 protein is 2 mg/mL. Considering that CAHS3 proteins are present only in the cytosol and not in the nucleus or extracellular space, the CAHS3 protein concentration is highly likely underestimated and the physiological concentration would be much higher, and we assumed it is not far from the concentration used in the gel transition experiments *in vitro*. Considering the cell volume reduction during dehydration which leads to a significant increase in both the protein concentration and ion strength that might be one of the gel-inducing factors as shown in Fig 7C, the intracellularly abundant CAHS proteins could undergo gel-transition in tardigrade cells and provide mechanical stabilization of cell integrity during dehydration (Fig 8). The cytoskeletal filamentous network formed by CAHS proteins under a stress condition may also support the cells to suppress harsh cell volume changes and deformation during water deficient stress (Fig 9). This gel-transition and/or cytoskeletal role could partly account for the exceptional stability of dehydrated tardigrades. The sol-gel transition and filament-formation of CAHS proteins were highly reversible and stress-dependent, and FRAP analyses revealed that CAHS proteins were immobile only when filaments formed under a stress condition. Therefore, we suppose that CAHS proteins are freely dispersed in a hydrated condition to minimize interference with other biological processes, whereas in a dehydrated condition, CAHS proteins form an intracellular filamentous network and elastic hydrogel to provide mechanical stabilization of cell integrity.

Although CAHS proteins exhibit no sequence similarity with any other cytoskeletal proteins, they formed cytoskeleton-like filamentous networks independently from the other cytoskeleton under a hyperosmotic stress (Fig 4 and S9 Fig) and were also functional in resisting the deformative mechanical forces in animal cells exposed to the water deficient stress (Fig 9). Hence, we propose CAHS proteins as a novel cytoskeletal protein family with stress-dependence and gel-forming ability. Although no known motifs are found in the primary sequence of CAHS proteins, the C-terminal region including the highly conserved CR1 and CR2 motifs was essential and sufficient for the filament formation (Fig 5). This region was mostly predicted as helical and to form a coiled-coil structure (Fig 6 and S13 Fig). This prediction was also supported by the previous circular dichroism (CD) spectroscopy of CAHS1 protein of *R. varieornatus,* another member of the CAHS family [15]. During the review of this manuscript, two related papers were published [34,35, which reported that two other CAHS proteins, i.e., CAHS1 *of R. varieornatus* and CAHS8 *of H. exemplaris,* formed fibrous structure and gel in a concentration-dependent manner *in vitro,* and the enriched helix structure in the C-terminal regions in either CAHS proteins were demonstrated by elaborate NMR analyses and/or CD spectroscopy under the condition forming filaments or gels. These recent structural analyses are in a good agreement with our structural predictions (Fig 5A and S13 Fig), although the necessity of such helix structure for filament/gel formation was not demonstrated. The severe impairments in the filament/gel formation by proline substitutions in either the CR1 or CR2 region (Fig 6 and 7E) indicate that the secondary structure of CR1 and CR2 plays important roles in CAHS filament/gel formation. Some intrinsically disordered proteins are reported to form a gel-like granule condensate via promiscuous binding through multivalent interaction sites [36], but in CAHS3 and CAHS12, single amino acid substitution is enough to disrupt both filament formation and gel transition, suggesting that the mechanism of filament/gel formation of CAHS proteins is likely not due to multivalent interactions, but rather to polymerization based on the secondary structure. The prediction of 3-dimentional structures by AlphaFold2 [37,38] suggested that CAHS3-min proteins form a helix in the CR1+CR2 region with high confidence (pLDDT = 70~90) and 2 CAHS3-min proteins form an anti-parallel dimer with the juxtaposition of each helical region where the charge and hydrophobicity distribution is consistent with the stabilization of 2 helix interactions (S21 Fig). This anti-parallel alignment is similar to the lamin tetramer formation [31], suggesting that the process of filament formation of CAHS proteins may be somewhat similar to intermediate filaments.

In contrast to filament-forming CAHS3 and CAHS12, CAHS8 alone formed granule-like condensates in both human and insect cells under a hyperosmotic condition (Fig 2A and S8 Fig). Recently, CAHS1 protein from *R. varieornatus* was also reported to form granules in response to hyperosmotic stress in human cultured cells [34], and these stress-dependent granule condensation by CAHS8 and CAHS1 resembled the stress-granule formation in mammalian cells that occurs through phase separation to create protective membrane-less compartments against stress [39,40]. A recent study revealed that another desiccation tolerance protein, AfrLEA6, which is a group 6 LEA protein of *Artemia franciscana,* also undergoes phase separation to form granules in insect cells [14] and protects enzyme activity from desiccation stress *in vitro* [41]. Like stress-granules and AfrLEA6 granules, CAHS8 granules exhibited certain sensitivity against 1,6-hexanediol treatment (Fig 2C). CAHS8 and CAHS1 might exert similar protective functions via granule condensation under stress conditions. Alternatively, in cells co-expressing CAHS8 and CAHS12, as shown in Fig 3, CAHS8 contributes to filament formation with CAHS12 in tardigrades.

Two well-known desiccation-tolerance protein families, LEA proteins and CAHS proteins, are mutually unrelated in the primary sequence, but both become helix-rich structure from unstructured state by dehydration [12,34]. TFE also induces similar conformational changes of both protein families [15,27] and thereby might mimic dehydration stress. The CAHS proteins which were identified through TFE-based screening, exhibited clear reversible condensation in animal cells in response to a hyperosmotic stress without TFE, suggesting that our isolation scheme (Fig 1A) successfully capture the reversibly condensing proteins under a water-deficient condition. In the DRYPs, stress-related unstructured proteins including CAHS and LEA proteins were enriched (Fig 1E and G), as well as translational proteins and cytoskeleton-related proteins (Fig 1D). These proteins might be incorporated into stress-dependent condensates like stress granules to be protected from stress. Alternatively, some of them like cytoskeletal proteins might be co-precipitated through entangling with CAHS filaments. Although CAHS proteins are conserved only in eutardigrades, proteins with similar properties might be present in other desiccation-tolerant organisms and may contribute to stress resistance. It is noteworthy that the related animal groups such as heterotardigrades or arthropods also lack the canonical cytoplasmic IFs but excellent anhydrobiotic ability is observed in some selected species such as *Echiniscus testudo* (a heterotardigrade), *Polypedilum vanderplanki* (a sleeping chironomid) and *Artemia* (a brine shrimp) [2,5,6]. These animals might possess another class of stress-dependent filament-forming proteins. Recently, a new heat-soluble protein family termed EtAHS was identified in *E. testudo* [20]. This protein family or other new ones are likely good candidates. Our isolation scheme of DRYPs may provide a general and potent method to identify unstructured proteins that undergo reversible condensation to filaments or granules in a stress-dependent manner from various organisms. CAHS proteins were originally identified by searching for heat-soluble proteins to identify anhydrobiotic protectants in tardigrades [15]. Later, many heat-soluble proteins were identified from humans and flies, dubbed Hero proteins [42], that exhibit no sequence similarity with CAHS proteins but provide stabilization of other proteins as CAHS and LEA proteins do. Similarly, future DRYPome analysis may lead to the identification of protective phase-separating proteins even in non-anhydrobiotic organisms.

In the present study, we established a new method to comprehensively identify proteins that are reversibly condensed in response to desolvating stress and found 336 such proteins from desiccation-tolerant tardigrades. The major components, CAHS3 and CAHS12, were shown to form cytoskeleton-like filaments and elastic hydrogel in a stress-dependent manner. Furthermore, we demonstrated that CAHS3 can confer mechanical resistance against deformation stress on insect cells and enhanced their tolerance to dehydration-like stress. We propose that these CAHS proteins function as novel stress-dependent and gel-forming cytoskeletal proteins that provide mechanical strength to stabilize cellular integrity during stress. Our data suggested a novel desiccation tolerance mechanism based on filament/gel formation. The isolation scheme established in this study opens the way to identifying such novel stress-dependent cytoskeletal proteins from various organisms.

## Materials and Methods

### Animals

We used the previously established YOKOZUNA-1 strain of the desiccation-tolerant tardigrade *R. varieornatus* reared on water-layered agar plate by feeding alga *Chlorella vulgaris* (Recenttec K. K., Japan) at 22°C as described previously [33].

### Identification of dehydration-dependent reversibly condensing proteins

Prior to protein extraction, tardigrades were starved for 1 day to eliminate digestive food. Approximately 400 *R. varieornatus* were collected and extensively washed with sterilized Milli-Q water to remove contaminants. Tardigrades were rinsed with lysis buffer, phosphate-buffered saline (PBS; 137 mM NaCl, 2.7 mM KCl, 10 mM Na_2_HPO_4_, 1.76 mM KH_2_PO_4_, pH 7.4) containing cOmplete protease inhibitors (Roche), and transferred to a 1.7 mL tube. Tardigrades were homogenized in 20 μL lysis buffer using a plastic pestle on ice. The pestle was rinsed with an additional 20 μL of lysis buffer collected in the same tube. After centrifugation at 16,000 × g for 20 min at 4°C, the supernatant was recovered as a soluble protein extract. To mimic dehydration stress, the desolvating agent, trifluoroethanol (TFE), was added (final concentration 10%, 20%, or 30%) and the mixture was incubated on ice for 1 h to allow complete induction of condensation. After centrifugation at 16,000 × g for 20 min, the supernatant was removed as a TFE-soluble fraction and the remaining precipitate was washed twice by lysis buffer containing TFE at the same concentration. The washed precipitate was resuspended in lysis buffer without TFE, and incubated at room temperature for 30 min to facilitate resolubilization. After centrifugation at 16,000 × g at 4°C, the supernatant was recovered as a resolubilized fraction. The fractions were analyzed by SDS-PAGE and proteins were visualized using a Silver Stain MS Kit (Fujifilm). Three selected bands were excised and separately subjected to mass spectrometry. Comprehensive identification of DRYPs was achieved by shot-gun proteomics of the resolubilized fraction. Briefly, proteins in gel slices or in the fraction were digested with trypsin and fragmented peptides were analyzed by nano LC-MS/MS. Proteins were identified using MASCOT software (Matrix Science). The mass spectrometry proteomics data have been deposited to the ProteomeXchange Consortium via the jPOST repository with the dataset identifier PXD030241.

### *In silico* structure predictions

The unstructured score of the proteins was calculated by IUPred2A [43]. IUPred2A produces the score for each amino acid position in a protein, and an average value was used as a score for each protein. A *de novo* protein sequence motif search in CAHS protein families was performed by the motif discovery tool, MEME version 5.0.4 [44] (https://meme-suite.org/meme/tools/meme). The parameters were as follows: (occurrence per sequence = 0 or 1; the maximum number to be found = 10; the motif width = 6 to 50). The secondary structures of CAHS3, CAHS8, and CAHS12 proteins were predicted by JPred4 [45] (https://www.compbio.dundee.ac.uk/jpred/). The coiled-coil regions of CAHS3 and CAHS12 were predicted by COILS [46] (https://embnet.vital-it.ch/software/COILS_form.html). The 3-dimensional structure prediction of the CAHS-min protein homo-dimer was performed by Alphafold2 [38] (https://colab.research.google.com/github/sokrypton/ColabFold/blob/main/AlphaFold2_complexes.ipynb). The 3-dimensional structures were visualized with UCSF ChimeraX v.1.2 [47].

### Enrichment analysis

To utilize well-annotated information in the model organism *Drosophila melanogaster,* we assigned a *D. melanogaster* ortholog for each *R*. *verieornatus* protein by a reciprocal BLAST search. We assigned 231 fly orthologs for 336 DRYPs and 7,361 fly orthologs for all 19,521 *R. varieornatus* proteins. Using the assigned fly orthologs, we performed enrichment analyses with PANTHER Overrepresentation Test [48] (PANTHER Protein Class version 16.0, Fisher’s test; http://pantherdb.org/) and Metascape [49] (GO Cellular Components; https://metascape.org/). The list of fly orthologs for all *R. varieornatus* proteins was used as a reference in the enrichment analyses.

Among tardigrade stress-related proteins described previously [16], 7 protein families containing more than 5 members were selected for the enrichment analysis. Enrichment of each family in DRYPs was statistically examined by Fisher’s exact test using R. Enrichment of up-regulated genes was similarly examined except a chi-square test was used.

### Differential gene expression analysis

Transcriptome data at a hydrated state and a dehydrated state were retrieved from the public database (DRR144971-DRR144973 and DRR144978-DRR144980 for *Paramacrobiotus* sp. TYO; SRR5218239-SRR5218241 and SRR5218242-SRR5218244 for *Hypsibius exemplaris,* respectively). The genome sequence of *Paramacrobiotus* sp. TYO was retrieved from the public database under accession numbers BHEN01000001-BHEN01000684 [11]. The genome sequence of *H. exemplaris* v3.0 was retrieved from http://www.tardigrades.org. RNA-seq reads were mapped to the genome sequence using HISAT2 v.2.1.0 [50]. Read counts for each gene region were quantified by featureCounts in SubRead package v.1.6.3 [51] and statistically compared by R package DESeq2 [52]. The genes with FDR < 0.01 were considered as differentially expressed genes. Orthologous gene relationships were determined by reciprocal BLAST searches among 3 tardigrade species.

### Cell lines

We obtained HEp-2 cells (RCB1889) from RIKEN BioResource Center (BRC). The identity of the cell line was validated by short tandem repeat profiling and the cell line was negative for mycoplasma contamination (RIKEN BRC). The cell was maintained in minimum essential medium (Nacalai Tesque) containing 10% fetal bovine serum (FBS, Cosmo Bio or BioWest) at 37°C, 5% CO_2_. *Drosophila* S2 cells (Gibco) were cultured at 28°C in Schneider’s *Drosophila* Medium (Gibco) supplemented with 10% heat-inactivated FBS (BioWest) and penicillin-streptomycin mixed solution (Nacalai Tesque).

### Plasmids

CAHS3, CAHS8, and CAHS12 coding sequences were amplified from the corresponding EST clones of *R*. *varieornatus* [16] and inserted into pAcGFP1-N1 or pAcGFP1-C1 (Clontech) with (GGGGS)3 linker using In-Fusion HD Cloning Kit (Takara). Plasmids to express CAHS deletion mutants (CAHS3ΔCtail, CAHS3ΔCR2-C, CAHS3ΔN-M2, CAHS3ΔN-M3, CAHS3-min, CAHS12ΔCtail, CAHS12ΔCR2-C, CAHS12ΔN-M1, CAHS12ΔN-H2, and CAHS12-min) or leucine-to-proline substitution mutants (CAHS3-L207P, CAHS3-L236P, CAHS3-L207P-L236P, CAHS12-L204P and CAHS12-L241P) were generated by inverse PCR and ligation, or PCR-based site directed mutagenesis. The CAHS3/8/12-mScarlet-I expression vector was generated from CAHS3/8/12-GFP expression vector by replacing AcGFP1 coding sequences with *mScarlet-I* sequence fragments [53] synthesized artificially (IDT). Expression constructs for various cytoskeleton or organelle marker proteins were obtained from Addgene (S1 Table). For bacterial expression of His6-tagged CAHS proteins, *CAHS3, CAHS8* or *CAHS12* coding sequences were amplified and inserted into pEThT vectors [15], and *CAHS3-GFP* was similarly inserted into a pCold-I vector (Takara). For expression in *Drosophila* cells, codon-optimized *CAHS3, CAHS8, CAHS12,* and *AcGFP1* DNA fragments were synthesized (Gene Universal) and inserted into pAc5.1/V5-His A vector (Invitrogen). The FUS-Venus plasmid was a kind gift from Dr. Tetsuro Hirose.

### Live cell imaging under hyperosmosis

We used HEp-2 cells for live-imaging of fluorescently-labeled proteins because HEp-2 cell were well sticky even under a stress condition and enabled precise inspections. HEp-2 cells were transiently transfected with an expression vector of fluorescently labeled proteins using Lipofectamine LTX & Plus Reagent (Invitrogen) for 48 h before stress exposure. Prior to microscopy, the medium was replaced with Hanks’ Balanced Salt Solution (HBSS) without the dications and phenol red. For exposure to hyperosmotic stress, the buffer was replaced with HBSS containing 0.4 M trehalose. The cells were stained with Hoechst 33342 (5 μg/mL, Lonza) to visualize nuclear DNA. Fluorescent signals were observed using a confocal microscope LSM710 (Carl Zeiss). The number of cells for each CAHS distribution pattern, such as dispersed, granules or filaments, were counted by 2 independent investigators and averaged counts were used. For time-lapse imaging in 3-dimensional space, we used the LSM-980 with Airyscan to perform super-resolution imaging. From the z-stack images, we generated orthogonal projections using ZEN 2.6 software. In time-lapse imaging experiments, a perfusion system KSX-Type1 (Tokai Hit) was used to replace the buffer. To visualize actin filaments by chemical staining, HEp-2 cells were treated with silicon-rhodamine dye probing actin (SiR-actin, Spirochrome) in HBSS containing the drug efflux inhibitor verapamil (10 μM, Tokyo Chemical Industry) for 2 h. For actin polymerization inhibition experiments, cells were treated with cytochalasin B (5 μM, Nacalai Tesque) for 60 min. Cells were then observed by a confocal microscope LSM-710 (Carl Zeiss).

### Fluorescence recovery after photobleaching (FRAP) analysis

HEp-2 cells were transiently transfected with the expression construct of CAHS3-GFP. The transfected cells were then exposed to isosmotic HBSS or hyperosmotic buffer, HBSS containing 0.4 M trehalose, to analyze the mobility of CAHS3-GFP in the dispersed or filament state, respectively. FRAP experiments were performed at room temperature using a confocal fluorescence microscope (FV1200, Olympus). A spot approximately 0.77 μm in diameter was photobleached at 100% laser power (wavelength 473 nm), and the fluorescence recovery curves were analyzed using the Diffusion Measurement Package software (Olympus). The fluorescence intensity was normalized by the initial intensity before photobleaching.

### Sensitivity to 1,6-hexanediol treatment

HEp-2 cells were transfected with expression vectors of CAHS3/8/12-AcGFP1 or FUS-Venus. After 48 h, cells were exposed on minimum essential medium supplemented with 0.4 M trehalose and 10% FBS for 1 h to induce the formation of granules or filaments. FUS protein was used as a control as it is known to be incorporated into liquid droplets under hyperosmosis [54]. After the addition of a liquid droplet disruptor, 1,6-hexanediol (final 5%), fluorescent images were captured at 0 and 30 min later by a confocal microscope LSM710 (Carl Zeiss). The fluorescence intensity was measured by Fiji and normalized to the initial fluorescence intensity of the granules or filaments.

### Immunofluorescence

HEp-2 cells expressing CAHS3 or CAHS3 mutants were exposed to HBSS containing 0.4 M trehalose for 60 min to induce filament-formation. The cells were then fixed in methanol at −30°C for 3 min and washed 3 times with PBS containing 0.1% Tween 20 (PBS-T). The cells were blocked with 2% normal goat serum (Abcam) for 1 h at room temperature and then reacted with 1/200 diluted antiserum against CAHS3 in 2% normal goat serum for 1 h at room temperature or 16 h at 4°C. The cells were washed 3 times with PBS-T, and then reacted with 1/1,000 diluted Alexa Fluor546 goat anti-guinea pig secondary antibody (Invitrogen) and 1 μg/mL DAPI in 2% normal goat serum for 1 h at room temperature. Fluorescent signals were observed using a confocal microscope LSM710 (Carl Zeiss).

### Protein preparation

Recombinant proteins were expressed as N-terminally His6-tagged proteins in *Escherichia coli* BL21(DE3) strains. CAHS3, CAHS8, and CAHS12 proteins were expressed using pET system (Novagen) essentially as described previously [15]. CAHS3-GFP and AcGFP1 were expressed using a cold shock expression system (Takara) essentially as described previously [55]. Bacterial pellets were lysed in PBS containing cOmplete EDTA-free protease inhibitors (Roche) by sonication. For CAHS3, CAHS8 and CAHS12, the supernatant was heated at 99°C for 15 min to retrieve heat-soluble CAHS proteins in a soluble fraction as described previously [15]. From the soluble fraction, His6-tagged proteins were purified with Ni-NTA His-Bind Superflow (Novagen) and dialyzed against PBS using a Pur-A-Lyzer™ Midi Dialysis Kit (Merck).

### *In vitro* polymerization of CAHS3-GFP proteins

Purified CAHS3-GFP or AcGFP1 protein solution in PBS (~ 40 μM) was directly dropped on cover glass (MATSUNAMI), and fluorescent images were captured by a confocal microscope LSM710 (Carl Zeiss). To induce the polymerization of CAHS3, an equal amount of PBS containing TFE was added (final 20%), and time-lapse images were captured every 5 s.

### *In vitro* gelation

Purified recombinant CAHS protein solution (5 mg/mL) was placed in a 0.2-mL tube. Inducing reagents such as TFE (final 20%), polyethylene glycol (final 20%), or NaCl (final 2 M) were added to the protein solution and incubated at room temperature for 10 min. Then, the tube contents were spread out on parafilm to check if it had solidified into a gel-like state or remained in a liquid state. Photos were obtained by a digital camera with a short focal length (Olympus TG-6).

### Preparation of cell-like microdroplets

Cell-like microdroplets coated with a lipid layer of phosphoethanolamine (Nacalai Tesque) were prepared in an oil phase. First, dry films of the lipids were formed at the bottom of a glass tube. Mineral oil (Nacalai Tesque) was then added to the lipid films followed by 90 min of sonication. The final concentration of the lipid/oil solution was approximately 1 mM. Next, 10 vol % of the protein solution (40 μM GFP-labeled CAHS3 or 40 μM GFP) was added to the lipid/oil solution at ~25°C. After emulsification via pipetting, the ~40 μL sample containing the microdroplets was placed on a glass-bottom dish. To condense the proteins inside the droplets upon dehydration, we added 40 μL salted oil. Mechanical measurements were performed 90 min after the droplet volume was approximately halved. For fluorescent imaging, 21 μM CAHS3-GFP and 171 μM CAHS3 were mixed and used.

### Measurement of the elasticity of droplets by micropipette aspiration

The elasticity of the cell-like microdroplets was evaluated by a micropipette aspiration system as reported previously [56]. The surface elasticity (Young’s modulus), *E*, is derived from the linear relationship between the elongation length into the micropipette, Δ*L,* and the aspiration pressure, Δ*P*: *E* = (3*ΔPRpΦ/2π)/ΔL,* wherein *R*p and Φ are the micropipette inner radius and wall function, which is derived from the shape of the micropipette. We used a micropipette with an *R*p smaller than × 0.4 of the microdroplet radius *R*. The value of Φ is 2.0. An increase in Δ*L* to above 50 μm under a very small Δ*P* (<< 0.5 kPa) indicates that the microdroplet is in liquid phase. In the case of the elastic gel phase, a linear relationship between Δ*L* and Δ*P* was confirmed for the small deformation within Δ*L* < 5 μm and Δ*P* < 3 kPa. Under these conditions, we derived the values of *E*. The temperature was approximately 25°C.

### Establishment of stably transfected cell line of *Drosophila* S2 cells

The expression vector Ac5-STABLE2-neo was obtained from Addgene (#32426) [57], and then the coding sequence of FLAG-mCherry was replaced with the codon-optimized CAHS3 coding sequence (Gene Universal) to express CAHS3-T2A-EGFP-T2A-neoR under the control of Ac5 promoter. The empty vector was constructed by deleting FLAG-mCherry from Ac5-STABLE2-neo, which was designed to express T2A-EGFP-T2A-neoR driven by the same Ac5 promoter. *Drosophila* S2 cells were transfected using a cationic liposome reagent Hilymax (Dojindo) with the expression construct or the empty vector above. We established stably transfected cells by culturing for 6 weeks under the drug selection with G418 disulfate (2000 μg/mL, Nacalai Tesque).

### AFM measurement of elasticity of S2 cells expressing CAHS3 protein

*Drosophila* S2 cells stably transfected with the CAHS3 expression construct and empty vector were cultured on PLL-coated coverslips (MATSUNAMI) at least one day before the measurement for cell attachment. For hyperosmotic treatment, the culture medium was replaced with the one containing 0.4 M trehalose 1 h before the measurement. The force spectroscopy was conducted using NanoWizard 3 Ultra AFM (Bruker) with an inverted microscope Olympus IX70 at 22°C. Cantilevers BL-AC40TS (Olympus) with a nominal spring constant of 0.09 N/m was calibrated using the thermal noise method for each experiment. Photodetector sensitivity was determined by fitting a line to the slope of the force distance curve acquired on the glass substrate. Indentation tests were performed at the speed of 2 μm/s for both approach and retraction, and the tests were repeated 16 times for each single cell. Young’s moduli of the cells were obtained by fitting the Hertz model to the force curves using indentation depths < 1 μm.

### Measurement of cell volume

The volume of each cell was measured using serial images of optical sections according to the previous publication [58]. Three-dimensional imaging was performed for GFP fluorescence in the stably transfected cells at 1.05 μm z-axis intervals using a 63×/1.2 oil-immersion lens on a confocal microscope LSM710 (Carl Zeiss). Cross-sectional area of the cell was calculated from each sectioned image using Fiji software and cell volume was estimated as a sum of them.

### Cell viability assay

As a hyperosmotic treatment, the S2 cells were exposed to the culture medium containing 0.4 M trehalose for 48 h. The cells were stained with Hoechst33342 (6.7 μg/mL, Lonza) and propidium iodide (PI, 0.67 μg/mL, Dojindo) for 30 min and observed with a fluorescence microscope BZ-X810 (Keyence). Hoechst33342-positive and PI-negative cells were counted as live cells and double-positive cells were counted as dead cells. The survival rates were calculated in 6 wells for each condition by counting 500-700 cells/well using analysis software Hybrid Cell Count (Keyence).

### Estimation of the amount of endogenous CAHS3 protein by immunoblotting

After extensive washing with purer water, approximately 100 *R. varieornatus* were lysed using pestle in 30 μL PBS containing cOmplete EDTA-free protease inhibitors (Roche) and centrifuged at 16,000 × g for 10 min. The soluble fractions of tardigrade lysate were mixed with 5 × SDS sample buffer (62.5 mM Tris-HCl pH6.8, 25% glycerol, 10% sodium dodecyl sulfate and 0.01% bromophenol blue) and 2-mercapto-ethanol. After heated at 100°C for 3 min, the samples were resolved by SDS-PAGE analysis and electroblotted onto PVDF membrane (Millipore). The membrane was blocked with 1% normal goat serum (Abcam) for 1 h at room temperature and reacted with the affinity-purified CAHS3 antibody diluted by 1% normal goat serum for 1 h at room temperature. After washed with TBS-T 3 times, the membrane was reacted with diluted peroxidase labeled anti-rabbit IgG antibody (KPL) for 1 h at room temperature. The membrane was washed with TBS-T 3 times, and then antibody-antigen complex was detected by ImageQuant LAS 500 (Cytiva) using enhanced chemiluminescence system (GE Healthcare). The diluted series of recombinant CAHS3 proteins (2.5, 5.0, 10.0, 20.0, 40.0 ng) were analyzed simultaneously on the same blot as quantification standards. Signal intensity of each corresponding band was measured by Fiji software and a linear regression was used to generate a standard curve between the signal intensity and the amount of protein as [Signal intensity] = [Amount of protein (ng)] × 446595.3952 - 696903.625; R^2^ = 0.9962. Using the well-fitted standard curve, the amount of endogenous CAHS3 protein was calculated to be ~3.81 ng per tardigrade.

## Acknowledgements

We are grateful to Tetsuro Hirose for providing the plasmid for FUS-Venus expression, and Tokiko Saigo for experimental assistance. Computations were partially performed on the NIG supercomputer at ROIS National Institute of Genetics.

## Supporting information

**S1 Movie. A three-dimensional image of CAHS3 filaments in a S2 cell.** Cytoskeleton-like distribution of CAHS3-GFP protein in *Drosophila* S2 cell under hyperosmotic cultured medium containing 0.4 M trehalose. Green indicates CAHS3-GFP and blue indicate Hoechst33342 staining of nuclei.

**S2 Movie. Movie of filament formation of CAHS3-GFP in HEp-2 cells.** Time after medium change to a hyperosmotic condition is shown. CAHS3-GFP simultaneously began to condense at many sites (155 s) and then elongated into filaments (235 s). Scale bar 5 μm.

**S3 Movie. Movie of CAHS3-GFP filament deformation in HEp-2 cells.** Time after hyperosmotic medium was replaced with isosmotic medium is shown. CAHS3-GFP filaments simultaneously collapsed and dispersed (400 s). Scale bar, 5 μm.

