## Supplemental Info for "Stress-dependent cell stiffening by tardigrade tolerance proteins through reversible formation of cytoskeleton-like filamentous network and gel-transition"

**S1 Table Fluorescent-tagged markers for various cytoskeletons and organelles used in this study**

| Addgene # | Plasmid/Marker name | Target structure | Fluorescent tag |
|---|---|---|---|
| 55065 | mCherry-keratin-17 | Keratin | mCherry |
| 55156 | mCherry-vimentin-7 | Vimentin | mCherry |
| 85047 | mScarlet-I-alpha-tublin-C1 | Tublin | mScarlet-I |
| 85056 | pLifeact-mScarlet-I | Actin | mScarlet-I |
| 98831 | mScarlet-I-lamin-b | Lamin | mScarlet-I |
| 85068 | pcytERM-mScarlet-I-N1 | Endoplasmic reticulum | mScarlet-I |
| 98818 | 4xmts-mScarlet-I | Mitochondria | mScarlet-I |

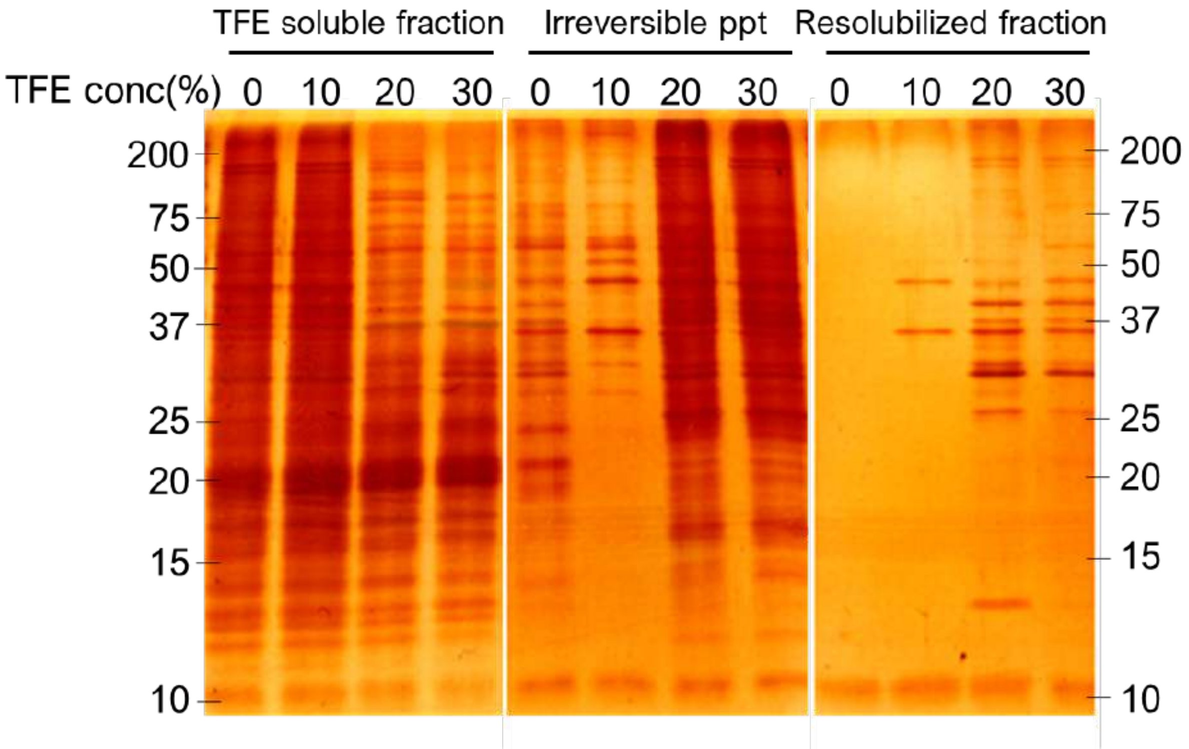


**S1 Fig. Silver-stained gel images of each fraction in the DRYP isolation process.** Each fraction was analyzed by SDS-PAGE and visualized by silver-staining. The image of the resolubilized fraction is partly presented in Fig 1B. As the concentration of TFE increased (0% to 20%), proteins decreased in the TFE soluble fraction, and proteins increased in both the irreversible precipitate (ppt) and the resolubilized fraction. Treatment with 20% and 30% TFE had largely similar effects.

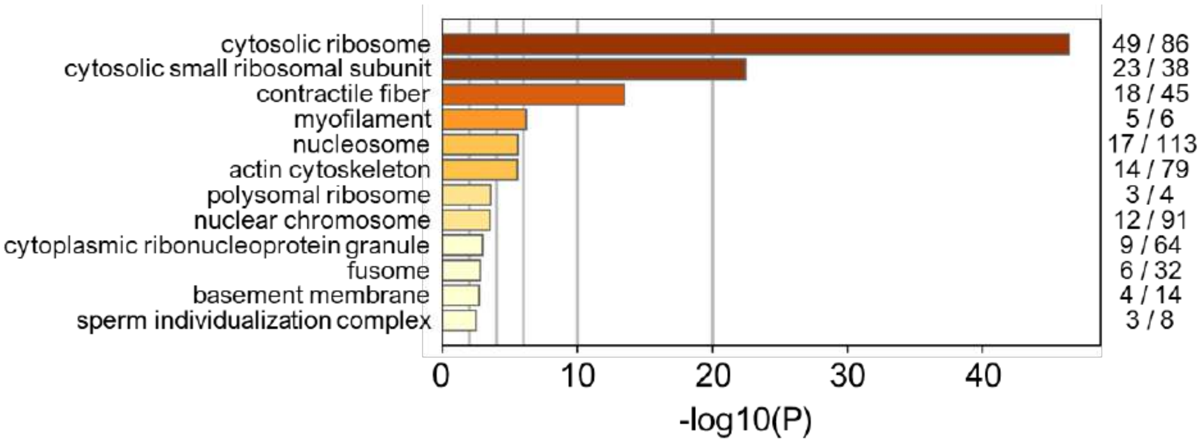


**S2 Fig. Enrichment analysis of Gene Ontology (GO) terms in DRYPs.** Ribosomal proteins and fiber proteins were highly enriched in DRYPs. GO terms of cellular components enriched in DRYPs were analyzed by Metascape.

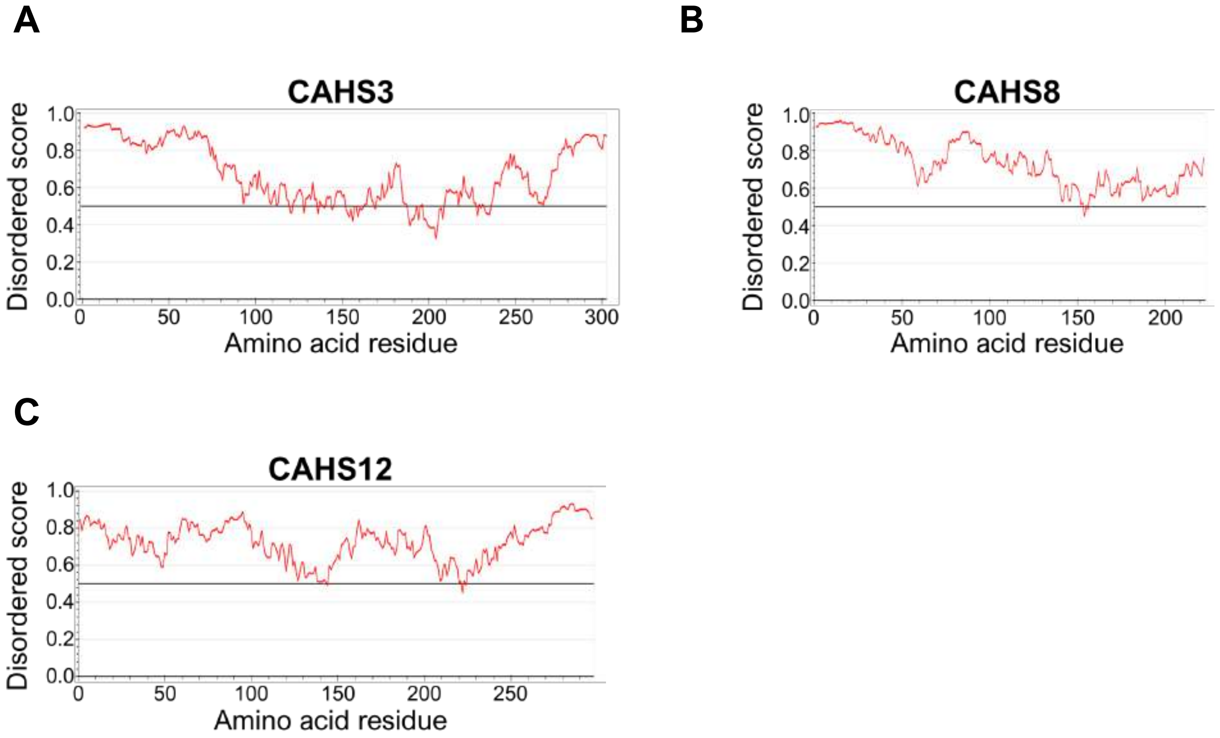


**S3 Fig. Prediction of disordered regions of CAHS3, CAHS8, and CAHS12 proteins.** (A–C) The unstructured score of each amino acid residue was calculated by IUPred2A for CAHS3 (A), CAHS8 (B), and CAHS12 (C). Scores above 0.5 indicate that the region is disordered. Each protein was predicted to be largely disordered throughout.

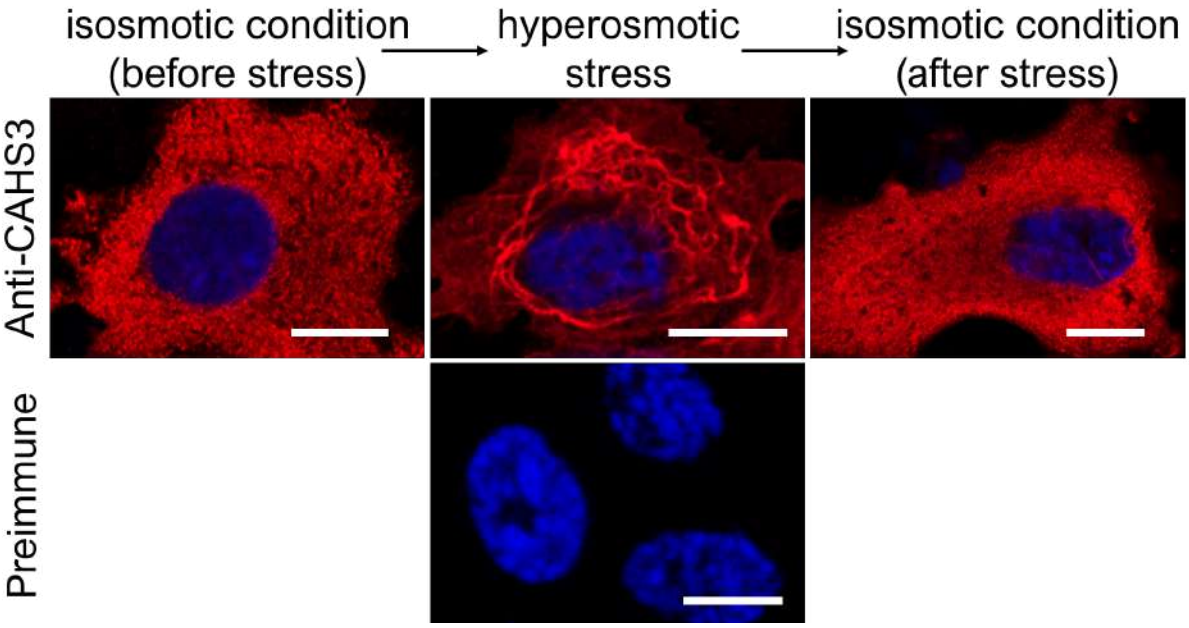


**S4 Fig. Hyperosmotic stress-dependent distribution changes in CAHS3 proteins without a GFP-tag in HEp-2 cells.** CAHS3 proteins were transiently expressed in HEp-2 cells and detected by immunofluorescence under isosmotic or hyperosmotic conditions. The detected distribution changes were similar to those of GFP-labeled CAHS3. Blue indicates DAPI staining of nuclei. Scale bar, 10 µm.

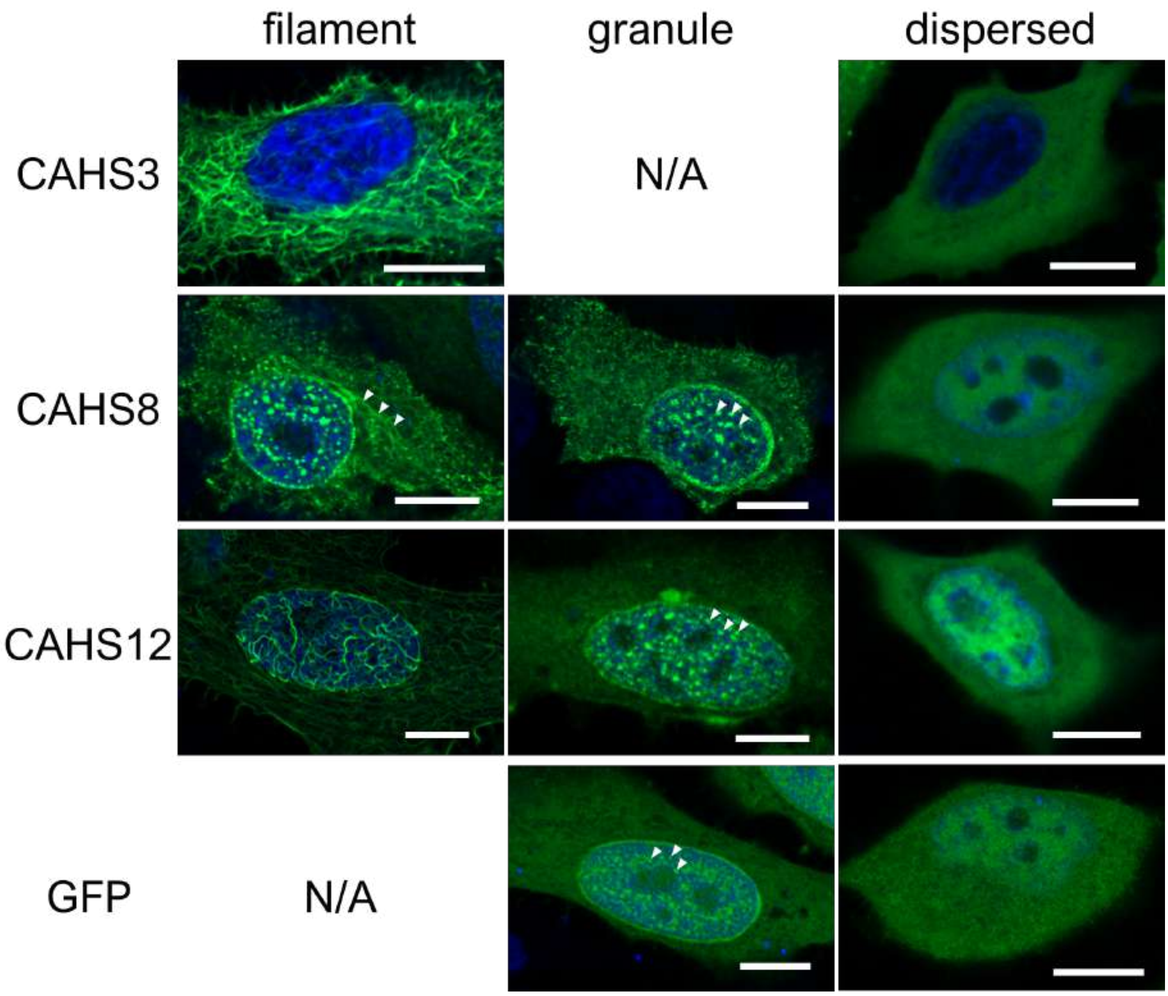


**S5 Fig. Representative images for each distribution pattern (filament, granule, or dispersed) of CAHS3-GFP, CAHS8-GFP, CAHS12-GFP, and GFP alone in human cultured HEp-2 cells.** N/A indicates that the corresponding distribution pattern is not or rarely found in a hyperosmotic condition. Scale bar, 10 µm.

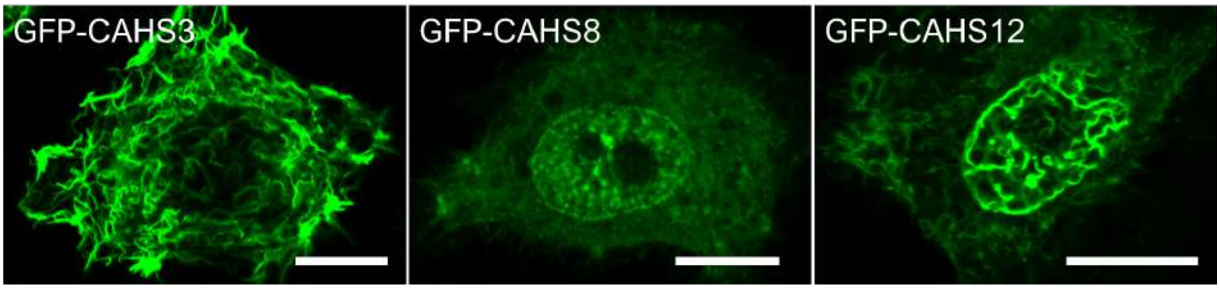


**S6 Fig. GFP-fusion to the other end (N-terminus) of CAHS proteins exhibited similar distribution patterns to those of C-terminally GFP-fused CAHS proteins under hyperosmosis.** N-terminally GFP-fused CAHS3 and CAHS12 exhibited filament-formation in response to hyperosmotic stress, and CAHS8 formed granule-like condensates like C-terminally GFP-fused CAHS proteins. The GFP-fusion site (N or C-terminus) did not affect the distribution pattern of CAHS proteins.

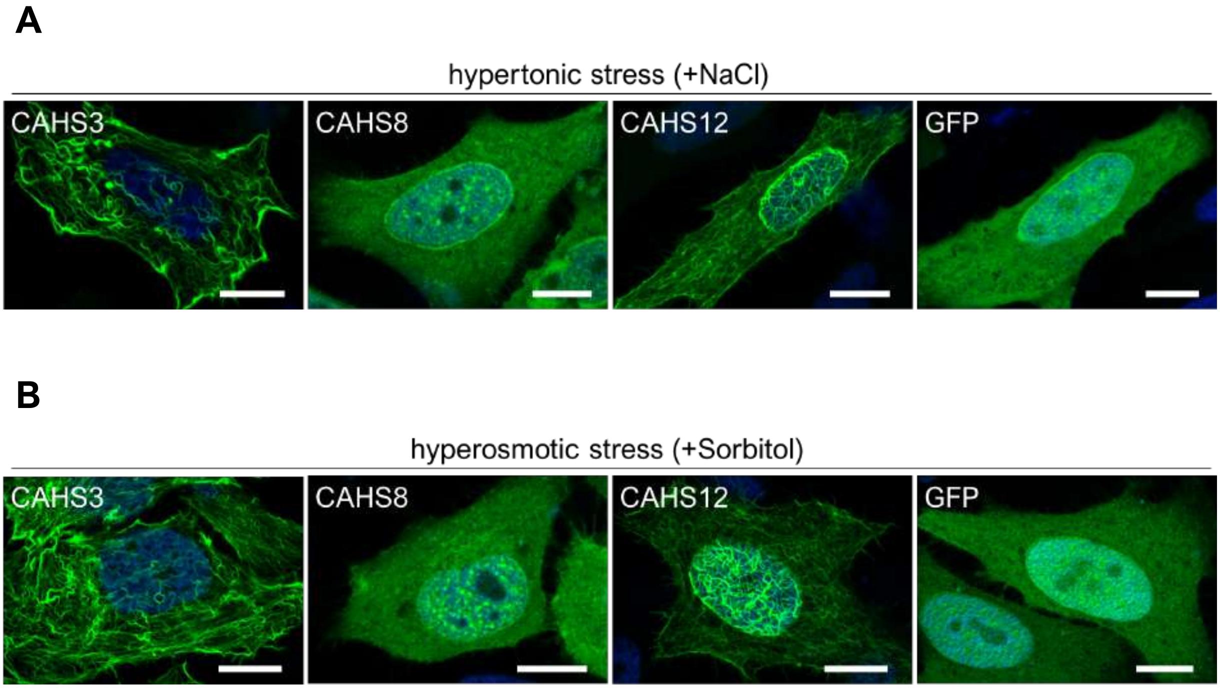


**S7 Fig. Effects of other hyperosmotic stressors (NaCl and sorbitol) on the distribution patterns of GFP-tagged CAHS proteins and GFP alone in human cultured cells.** (A–B) Representative distribution patterns of GFP-tagged CAHS proteins and GFP alone under hypertonic medium supplemented with 0.2 M NaCl (A) or hyperosmotic medium supplemented with 0.4 M sorbitol (B). Distribution changes were similar to those observed when treated with 0.4 M trehalose (Fig 2A). Blue indicates Hoechst33342 staining of nuclei. Scale bar, 10 µm.

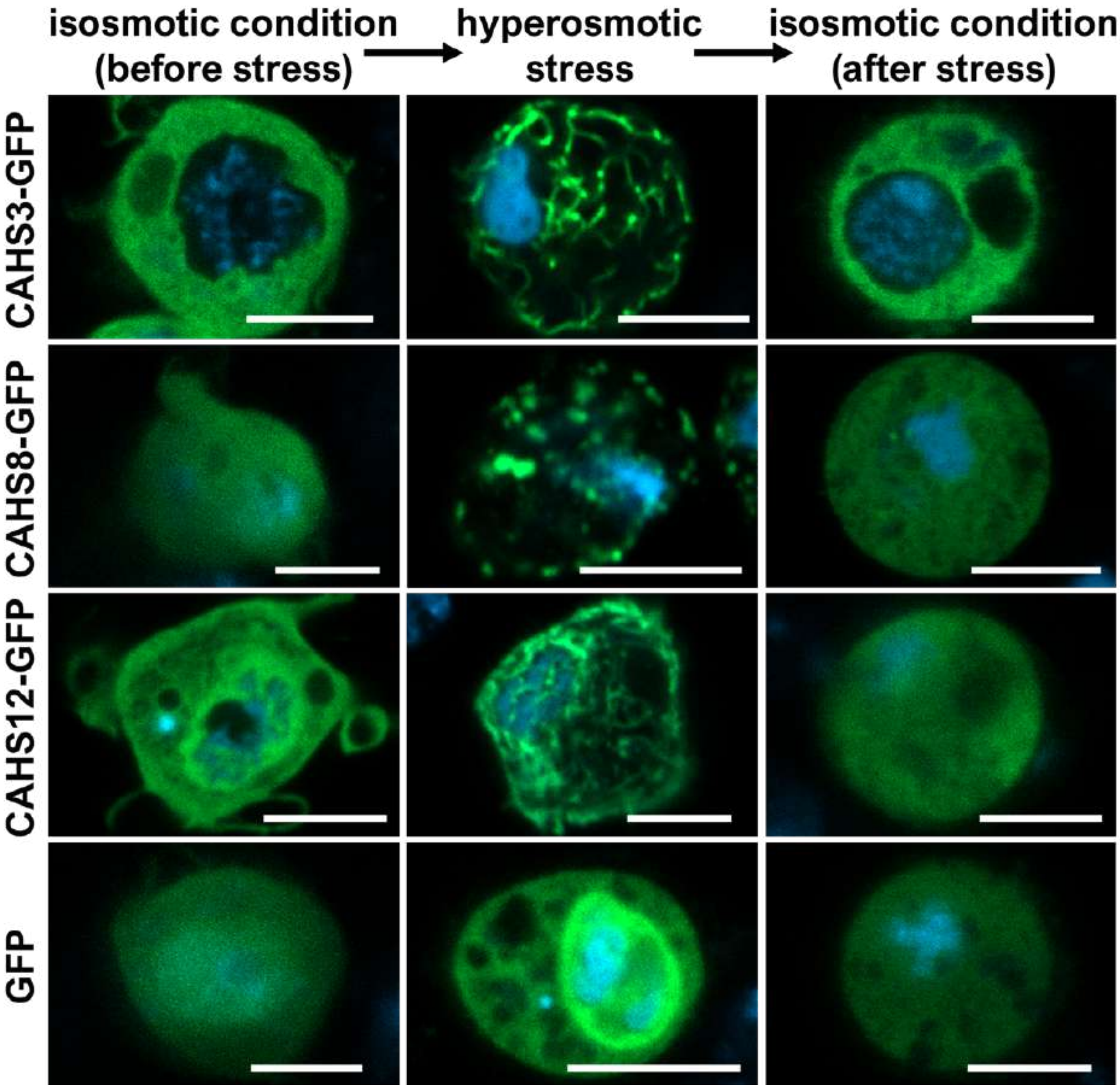


**S8 Fig. Distribution changes of CAHS-GFP proteins in *Drosophila* cultured S2 cells during transient hyperosmotic treatment.** Like in human cells, CAHS3-GFP and CAHS12-GFP reversibly formed filaments and CAHS8-GFP reversibly formed granules upon hyperosmotic stress in fly cells. As a hyperosmotic medium, the culture medium containing 0.4 M trehalose was used. Blue indicates Hoechst33342 staining of nuclei. Scale bar, 5 µm.

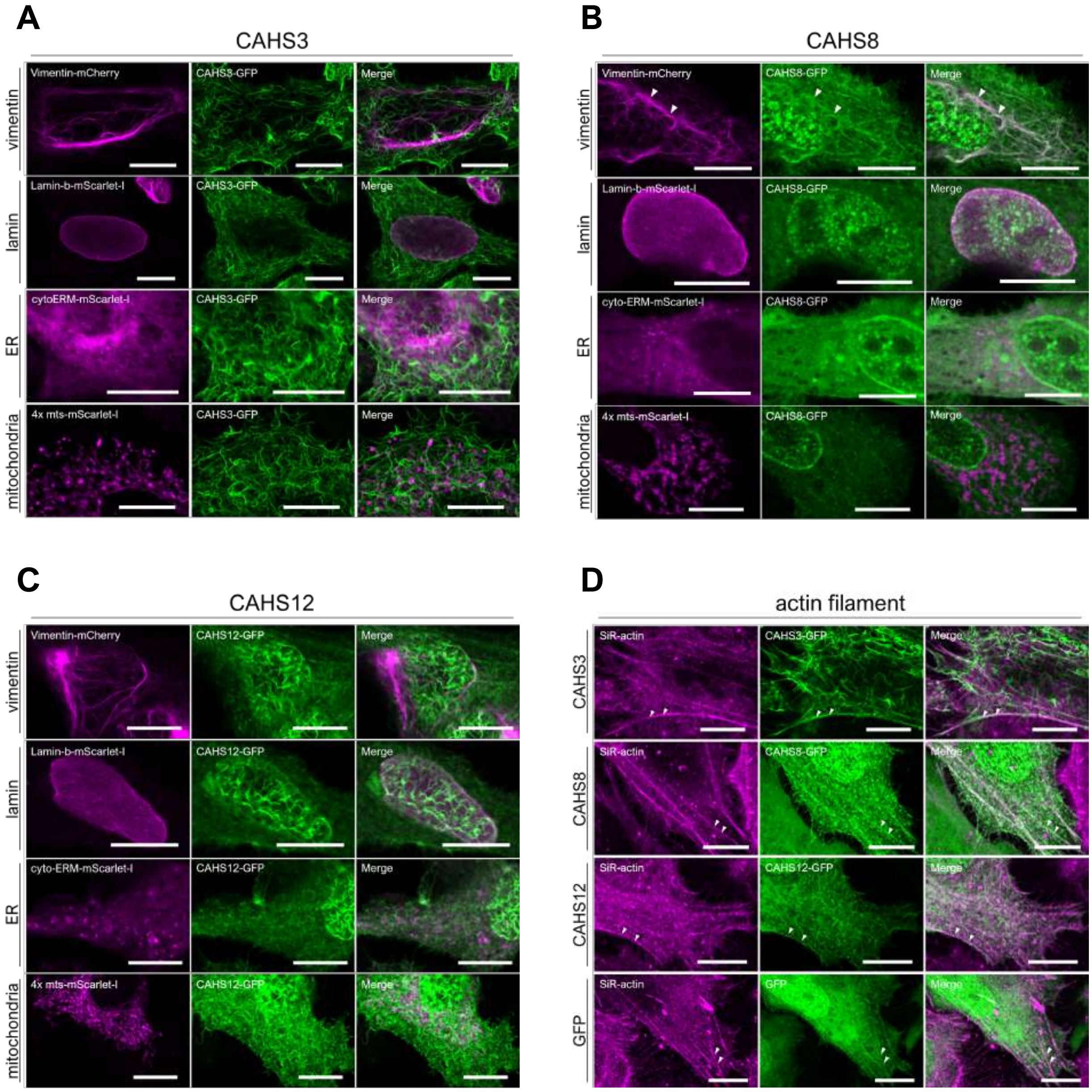


**S9 Fig. Co-localization analyses between CAHS proteins and cytoskeletons or organelles.** (A–C) Confocal images of HEp-2 cells expressing AcGFP1-tagged CAHS3 (A), CAHS8 (B) or CAHS12 (C) and other fluorescently labeled intermediate filaments (vimentin and keratin) or organelle markers (endoplasmic reticulum and mitochondria) under hyperosmosis. White arrows indicate detected co-localization. (D) Co-localization analyses between intrinsic actin filaments and CAHS-GFP proteins or GFP alone. Actin filaments was visualized by staining with the chemical probe SiR-actin. All examined proteins including GFP alone slightly co-localized with actin filaments similarly to the data visualized by Lifeact-mScarlet-I (Fig 4). Scale bar, 10 µm.

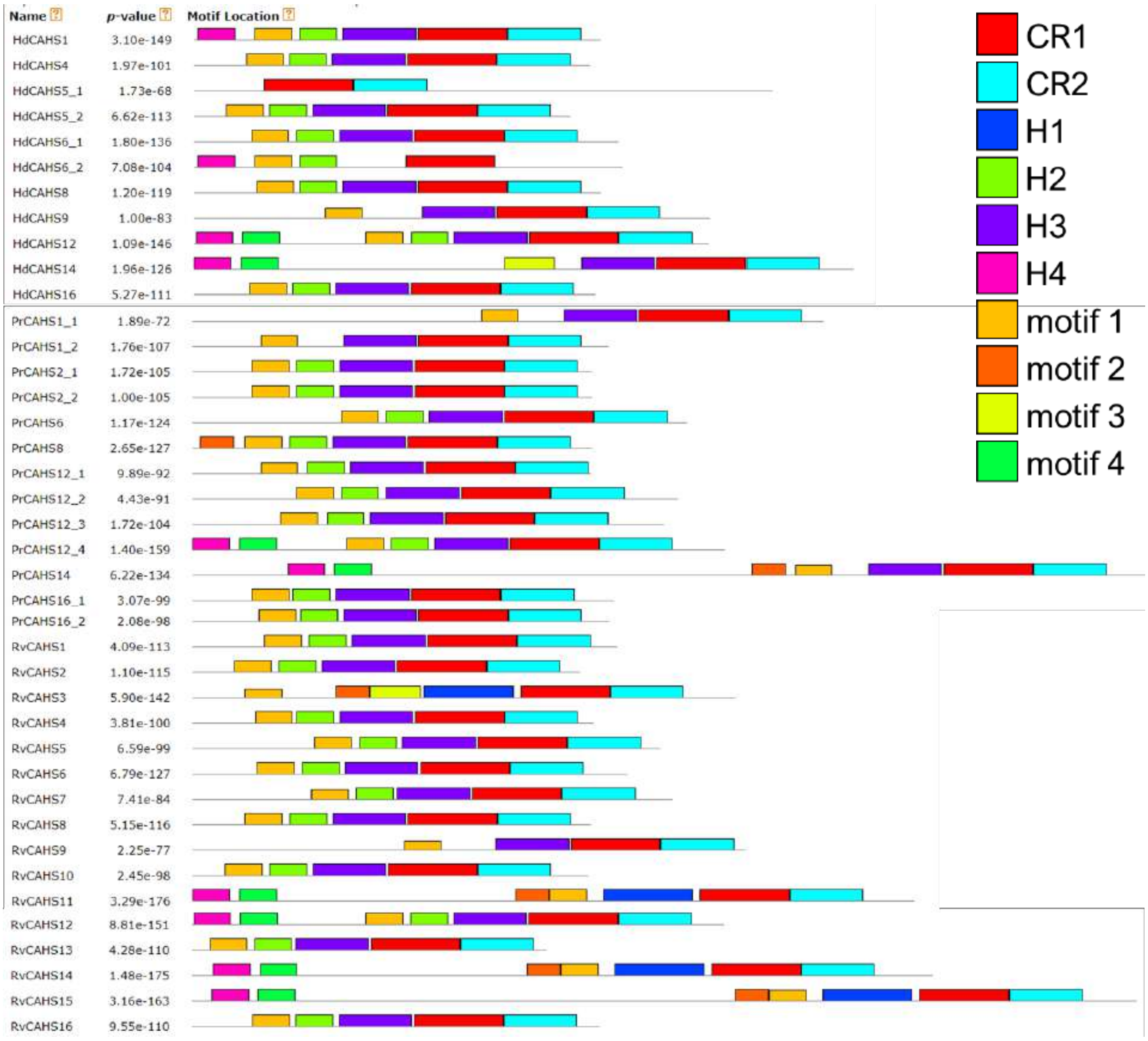


**S10 Fig. Informatically extracted motif structures of CAHS protein family.** Ten conserved sequence motifs were identified by MEME among 40 CAHS proteins from 3 tolerant tardigrades (*Hypsibius exemplaris*, *Paramacrobiotus* sp. TYO, and *Ramazzottius varieornatus*). Each motif is shown in the corresponding colored box. Both CR1 and CR2 were conserved in all 40 CAHS proteins except CR2 in HdCAHS6_2. CR1, CR2, H1, H2, H3, and H4 were predicted as helical regions by JPred4 (S12 Fig). Hd, *H. exemplaris* (formerly *H. dujardini*); Pr, *Paramacrobiotus* sp. TYO; Rv, *R. varieornatus*.

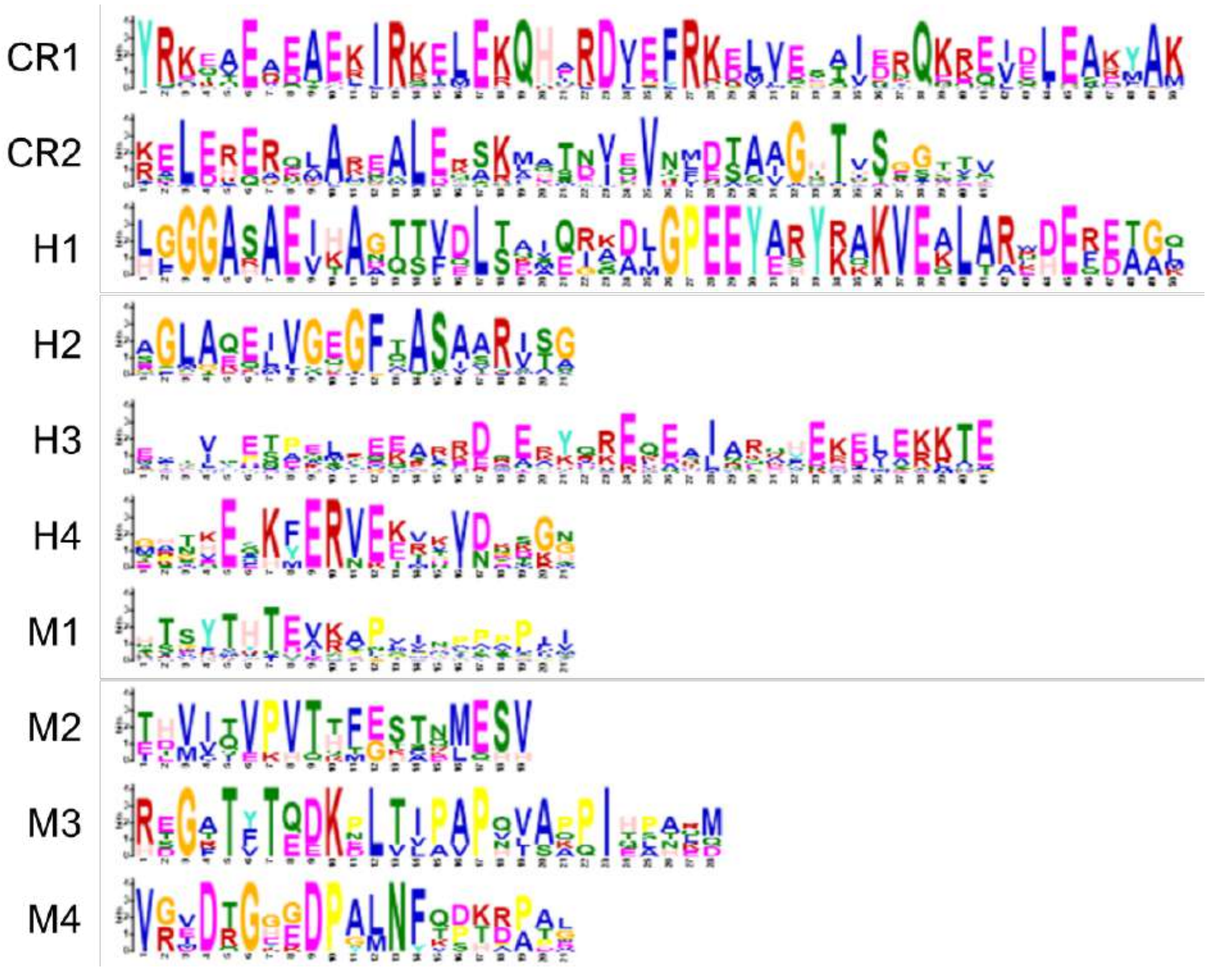


**S11 Fig. Sequence logo representation of the conserved protein sequence motifs among the CAHS protein family.**

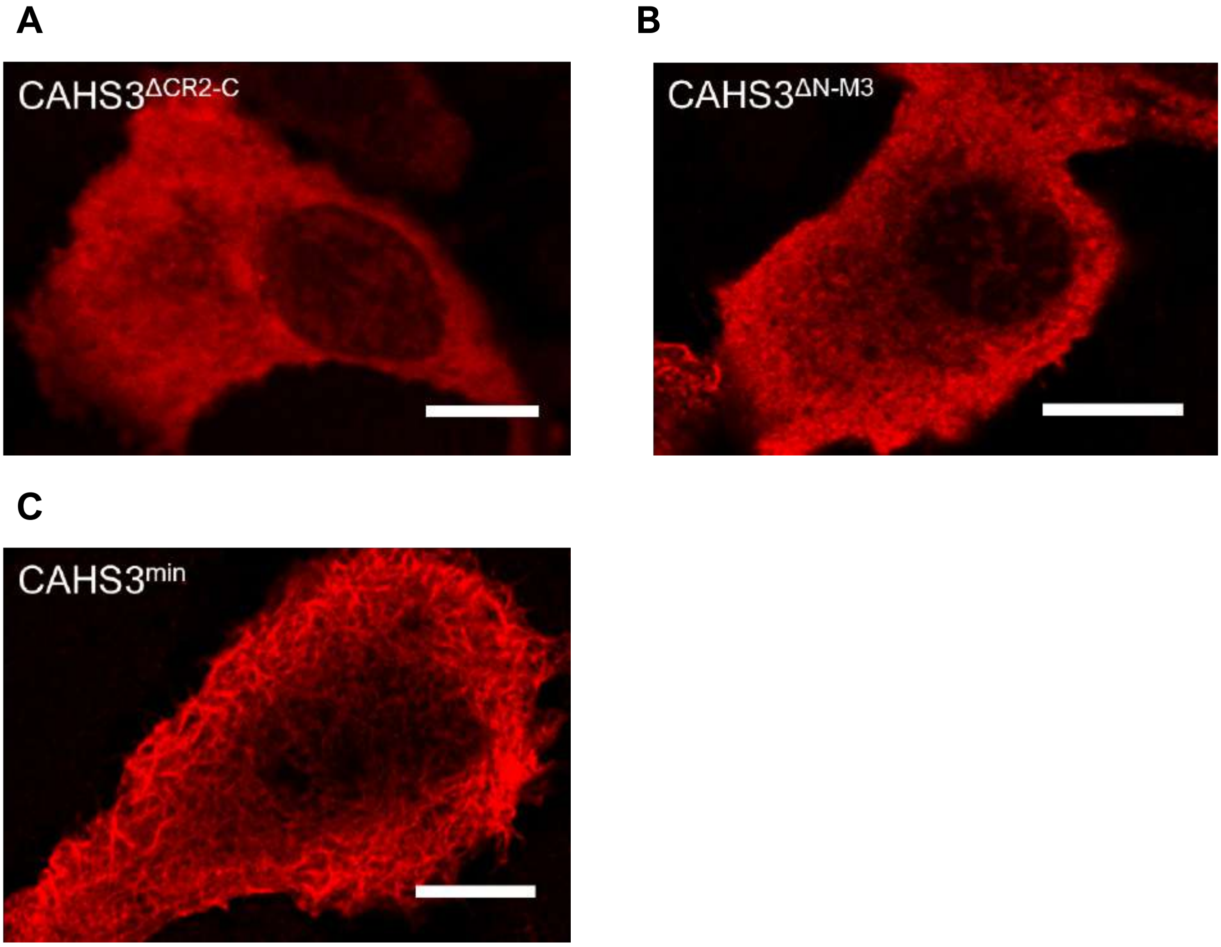


**S12 Fig. Distribution patterns of CAHS3 truncated mutants without GFP-tag in HEp-2 cells under a hyperosmotic condition.** (A–C) CAHS truncated mutants were expressed in HEp-2 cells and their distribution patterns were detected by immunofluorescence under a hyperosmotic condition. CAHS3ΔCR2-C (A) and CAHS3ΔN-M3 (B) failed to form long filamentous networks, whereas CAHS3-min (C) successfully formed filaments. The detected distribution patterns were similar to those of the corresponding CAHS3 mutants labeled with GFP (Fig 5B). Scale bar, 10 µm.

**A**

CAHS3

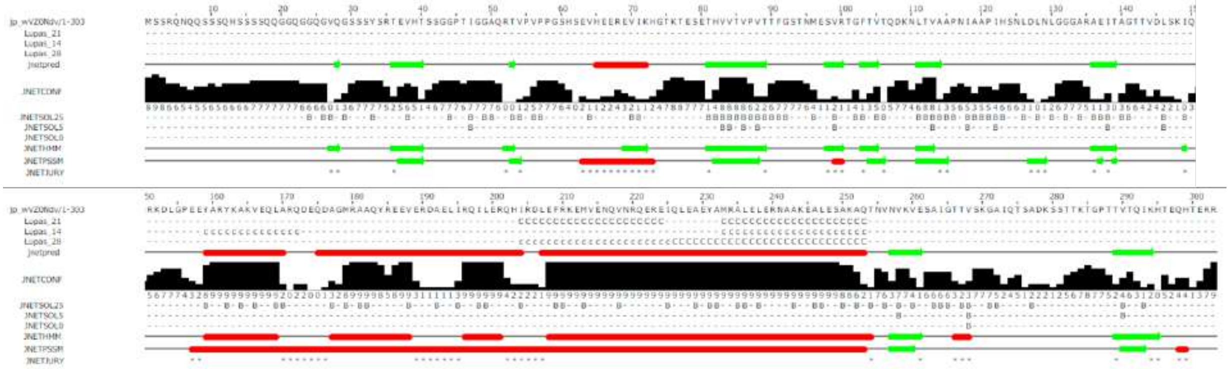

**B**

CAHS8

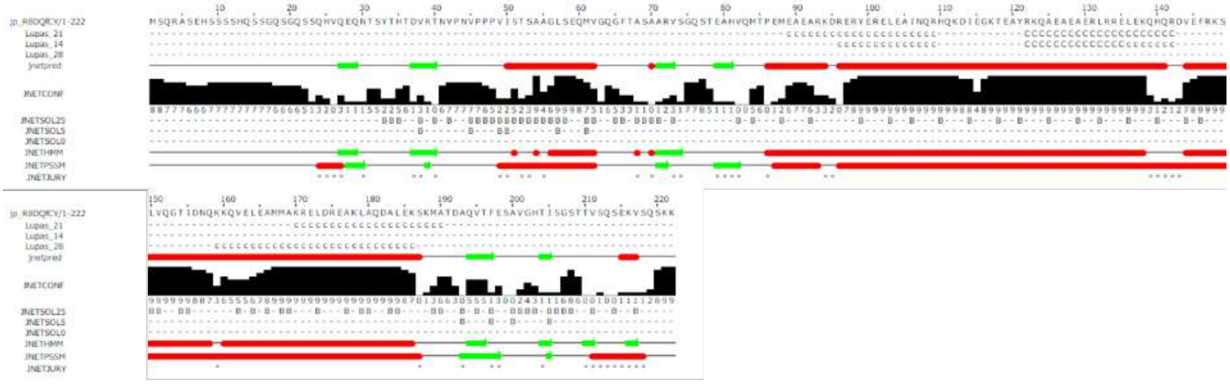

**C**

CAHS12

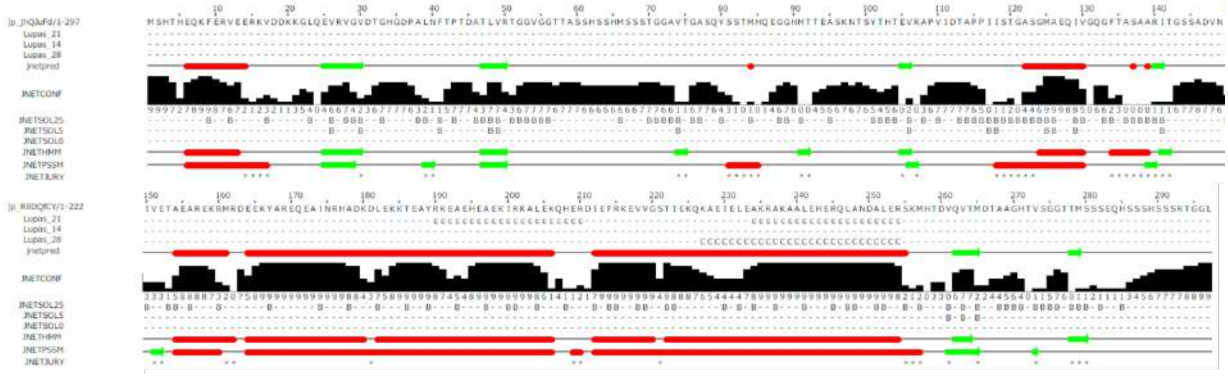

**S13 Fig. Predicted secondary structures of CAHS proteins.** Secondary structure predictions by JPred4 are shown for CAHS3 (A), CAHS8 (B), and CAHS12 (C). Red boxes indicate putative helical regions and green arrows indicate putative beta sheet regions in jnetpred, JNETHSSM and JNETPSSM respectively. Lupas shows coiled-coil prediction; ‘C’ or ‘c’ indicate putative coiled-coil region and the capital ‘C’ indicates a higher probability. JNETSOL show solvent accessibility.

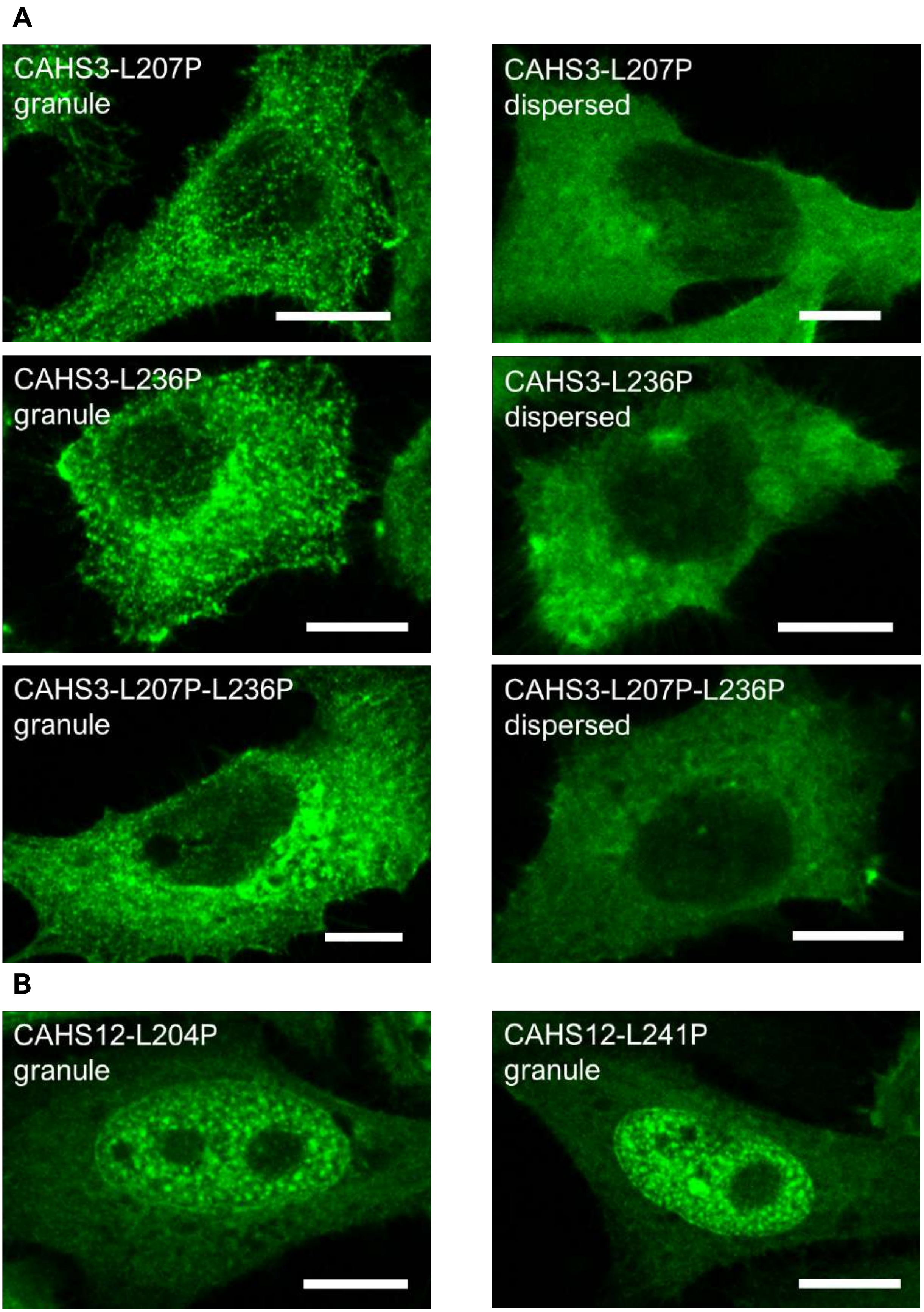


**S14 Fig. Representative images of each distribution pattern of proline-substituted CAHS3 and CAHS12 mutants.** (A–B) Representative images of granule-like condensation or dispersed distribution of proline-substituted mutants of CAHS3 (A) and CAHS12 (B). Scale bar, 10 μm.

A

B

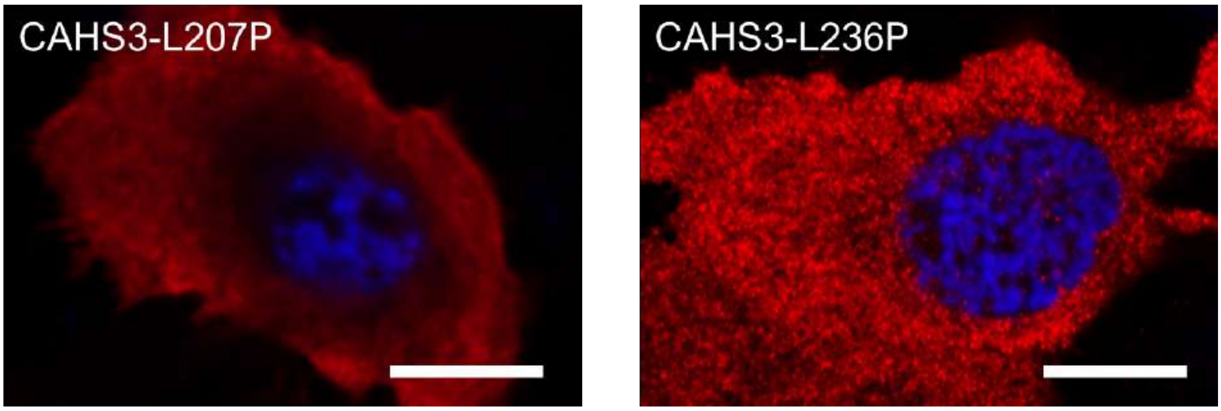


**S15 Fig. Distribution patterns of proline-substituted CAHS3 mutants without a GFP-tag in HEp-2 cells under a hyperosmotic condition.** (A–B) Distribution patterns of proline-substituted CAHS3 mutants were examined by immunofluorescence for both CAHS3-L207P (A) and CAHS3-L236P (B). Immunostaining images show the dispersed distribution or slightly condensed granules similar to the corresponding CAHS3 mutants labeled with GFP (Fig 6). Blue indicates DAPI staining of nuclei. Scale bar, 10 µm.

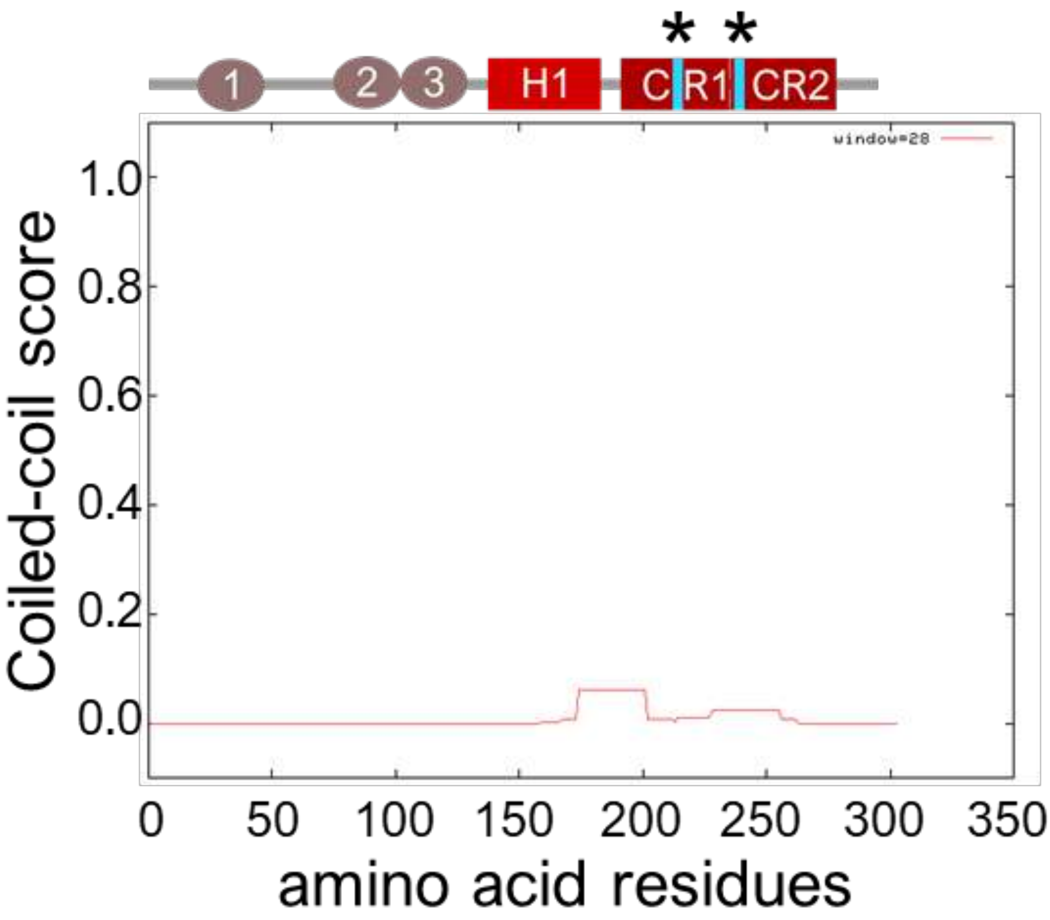


**S16 Fig. Significant decrease of the coiled-coil score in the double proline-substituted CAHS3 mutant.** Asterisks indicate the proline-substituted mutation sites. Coiled-coil score predicted by COILS is shown for CAHS3-L207P-L236P.

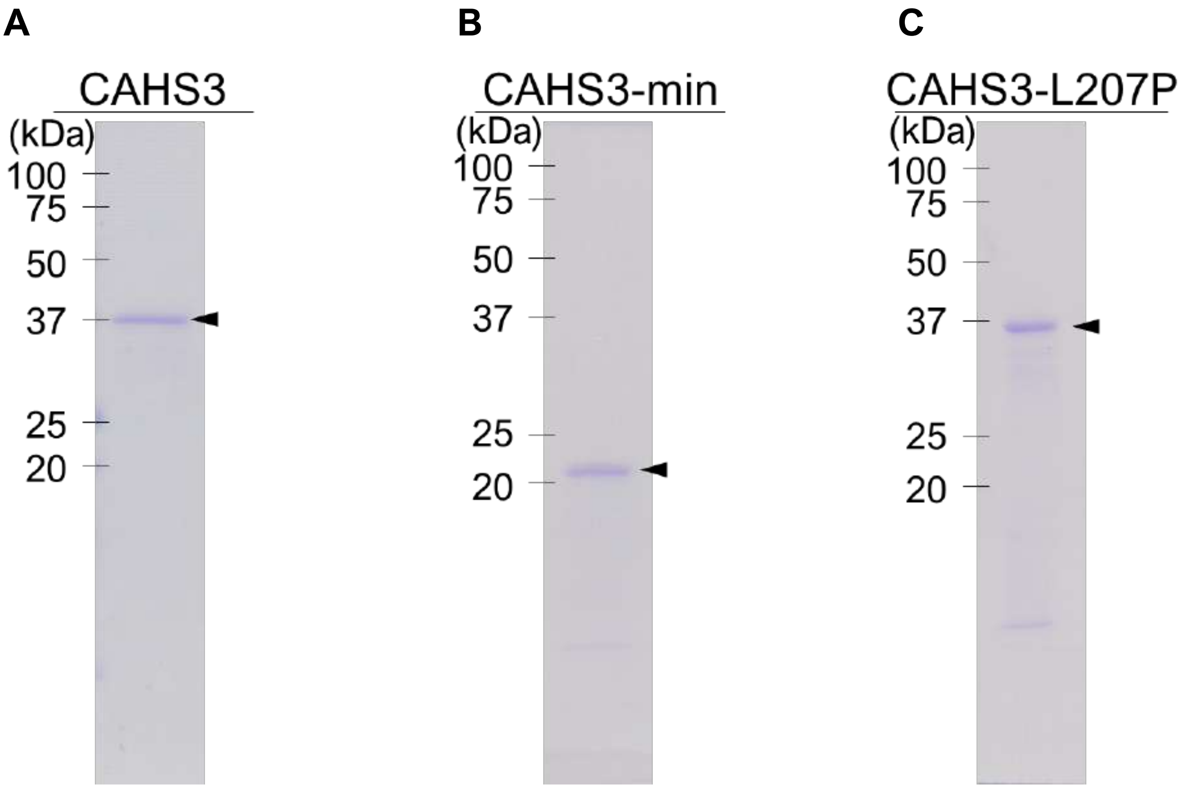


**S17 Fig. SDS-PAGE gel images of purified full-length CAHS3 and mutant recombinant proteins.** (A–C) Arrowheads indicate major bands corresponding to the expected length of full-length CAHS3 (A), CAHS3-min (B) and CAHS3-L207P proteins (C).

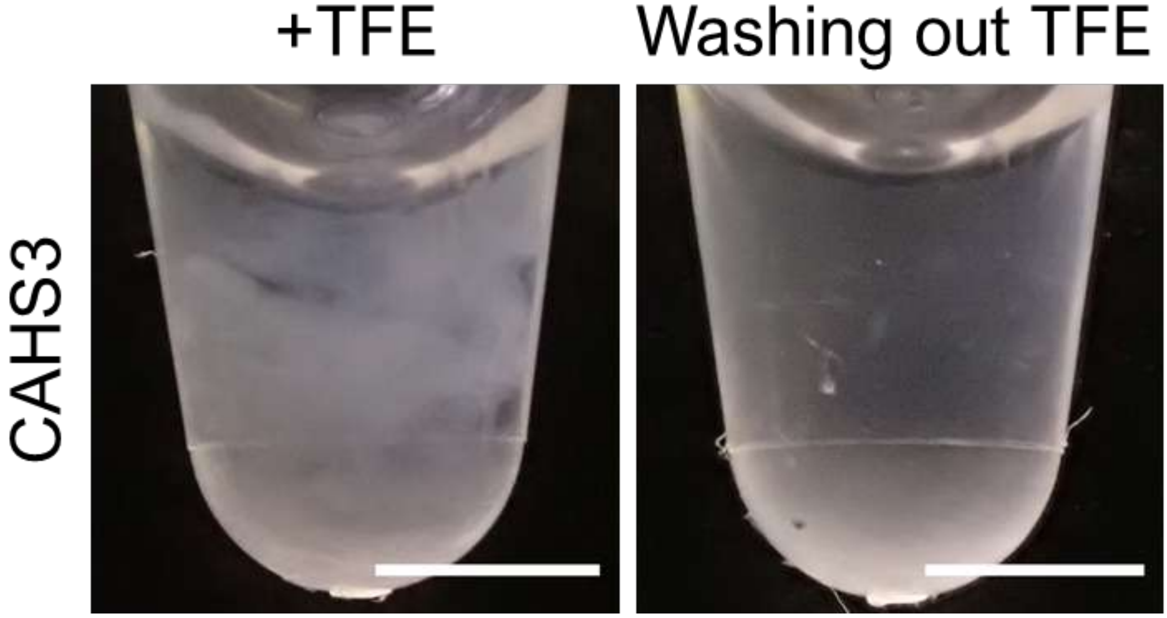


**S18 Fig. Resolubilization of TFE-dependent CAHS3 gelation.** CAHS3 gel condensates induced by TFE (final 20%) were redissolved by rinsing with TFE-free PBS. Scale bar, 2 mm.

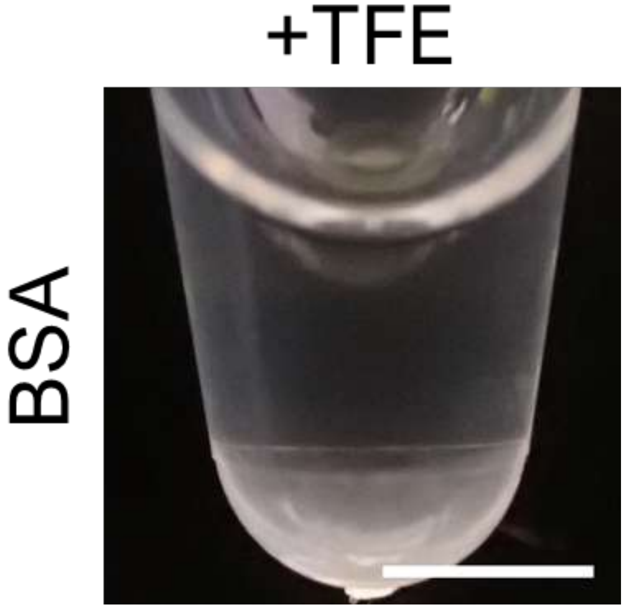


**S19 Fig. Effect of TFE on BSA solution.** TFE (final 20%) had no visible effect on BSA solution (final 4.0 mg/mL). Scale bar, 2 mm.

**A**

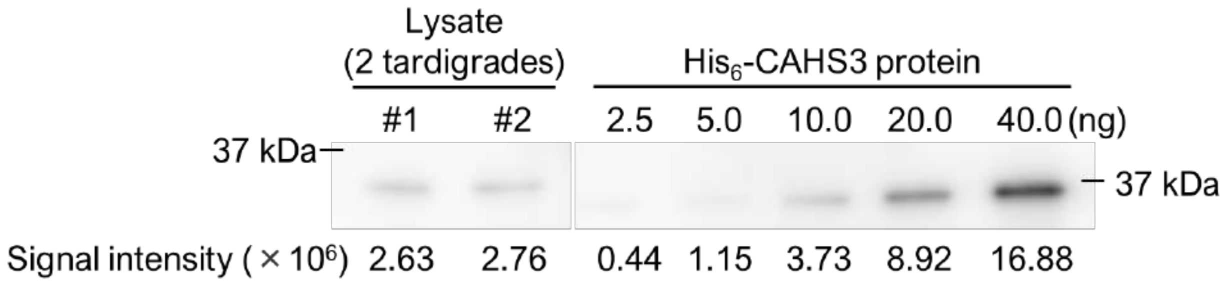


**B**

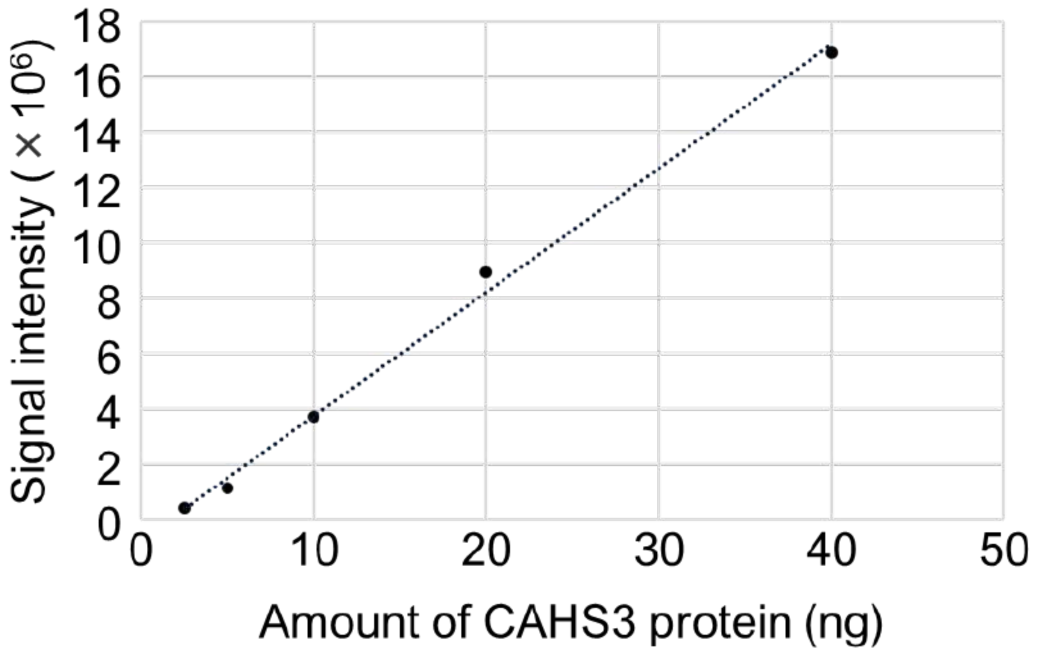


**S20 Fig. Estimation of the amount of CAHS3 protein in tardigrades.** (A) Endogenous CAHS3 protein in *R. varieornatus* lysate was detected by immunoblotting using anti-CAHS3 antibody. Each lane of the lysate (#1 and #2) contains protein amount corresponding to two individuals. Diluted series of recombinant CAHS3 proteins were simultaneously analyzed on the same blot as quantification standards. Due to additional $His_6$-tag, recombinant CAHS3 proteins exhibited slightly higher molecular weight than endogenous ones. Signal intensities were quantified using Fiji imaging software. (B) Based on the signal intensity of quantification standards, the standard curve was generated by a linear regression ($R^2$ = 0.9962). The amount of endogenous CAHS3 protein was estimated as about 3.81 ng per tardigrade.

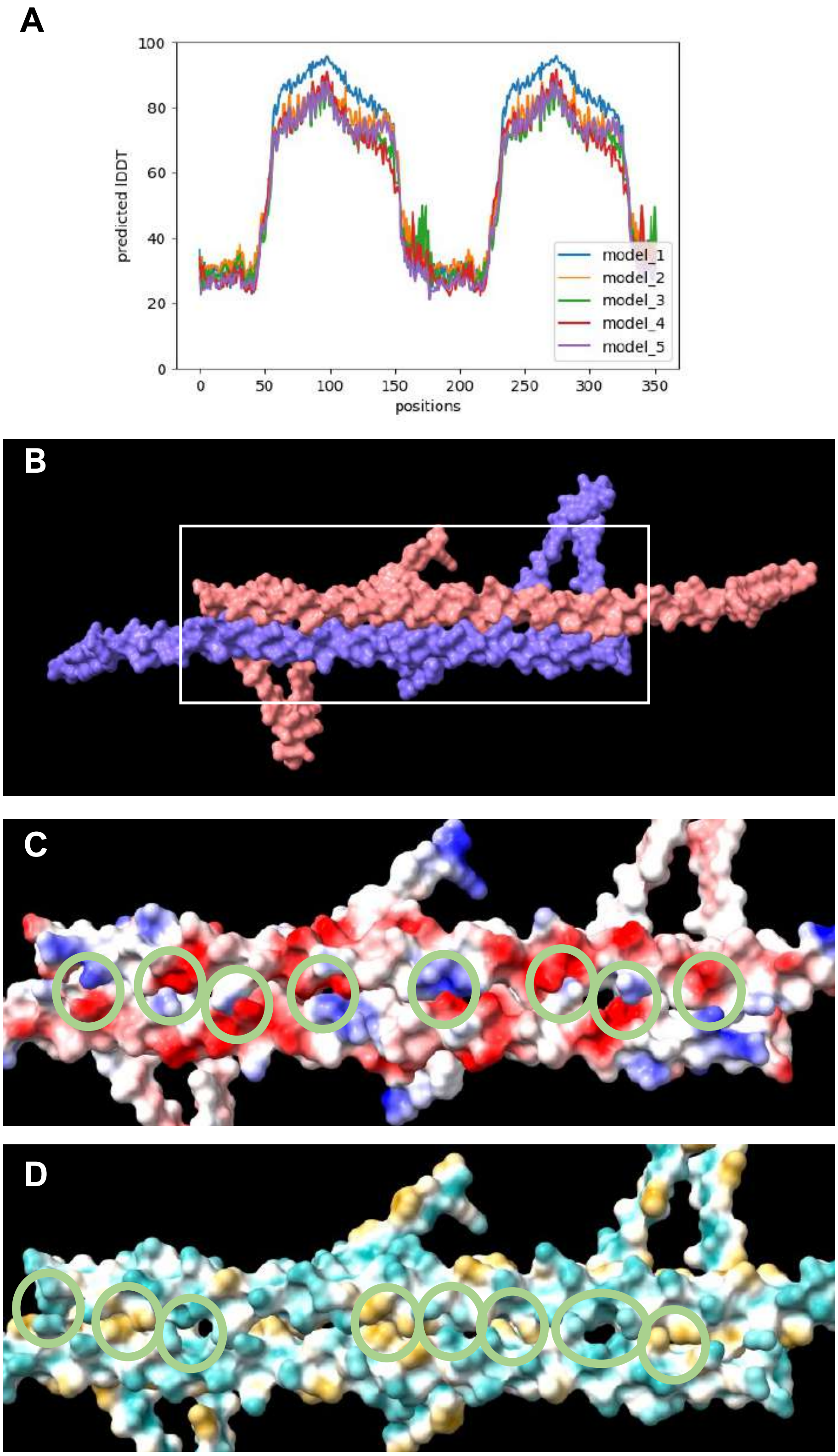


**S21 Fig. Predicted 3-dimensional structures of a homo-dimer of CAHS3-min proteins by AlphaFold2.** (A) pLDDT scores on the prediction corresponding to a tandem CAHS-min amino acid sequence. Scores corresponding to CR1+CR2 regions (70~90) indicated high structure confidence. (B) Two chains of CAHS3-min proteins distinguished by 2 colors. White box indicates the anti-parallel helical region. (C) Magnified view of the charge distributions in the juxtaposed helical regions. Green circles indicate the facing of opposite charges between 2 CAHS3-min proteins, suggesting stabilization by electrostatic interactions. (D) Magnified view of the hydrophobicity distributions. Green circles indicate the juxtaposition of similar hydrophobicities/hydrophilicities between 2 proteins, supporting hydrophobic interactions.
